# Hypoxia-inducible factor 2 is a key determinant of manganese excess and polycythemia in SLC30A10 deficiency

**DOI:** 10.1101/2023.02.20.529270

**Authors:** Milankumar Prajapati, Jared Z. Zhang, Courtney J. Mercadante, Heather L. Kowalski, Bradley Delaney, Jessica A. Anderson, Shuling Guo, Mariam Aghajan, Thomas B. Bartnikas

## Abstract

Manganese is an essential yet potentially toxic metal. Initially reported in 2012, mutations in SLC30A10 are the first known inherited cause of manganese excess. SLC30A10 is an apical membrane transport protein that exports manganese from hepatocytes into bile and from enterocytes into the lumen of the gastrointestinal tract. SLC30A10 deficiency results in impaired gastrointestinal manganese excretion, leading to severe manganese excess, neurologic deficits, liver cirrhosis, polycythemia, and erythropoietin excess. Neurologic and liver disease are attributed to manganese toxicity. Polycythemia is attributed to erythropoietin excess, but the basis of erythropoietin excess in SLC30A10 deficiency has yet to be established. Here we demonstrate that erythropoietin expression is increased in liver but decreased in kidneys in Slc30a10-deficient mice. Using pharmacologic and genetic approaches, we show that liver expression of hypoxia-inducible factor 2 (Hif2), a transcription factor that mediates the cellular response to hypoxia, is essential for erythropoietin excess and polycythemia in Slc30a10-deficient mice, while hypoxia-inducible factor 1 (HIF1) plays no discernible role. RNA-seq analysis determined that Slc30a10-deficient livers exhibit aberrant expression of a large number of genes, most of which align with cell cycle and metabolic processes, while hepatic Hif2 deficiency attenuates differential expression of half of these genes in mutant mice. One such gene downregulated in Slc30a10-deficient mice in a Hif2-dependent manner is hepcidin, a hormonal inhibitor of dietary iron absorption. Our analyses indicate that hepcidin downregulation serves to increase iron absorption to meet the demands of erythropoiesis driven by erythropoietin excess. Finally, we also observed that hepatic Hif2 deficiency attenuates tissue manganese excess, although the underlying cause of this observation is not clear at this time. Overall, our results indicate that HIF2 is a key determinant of pathophysiology in SLC30A10 deficiency.

**Graphical abstract:** 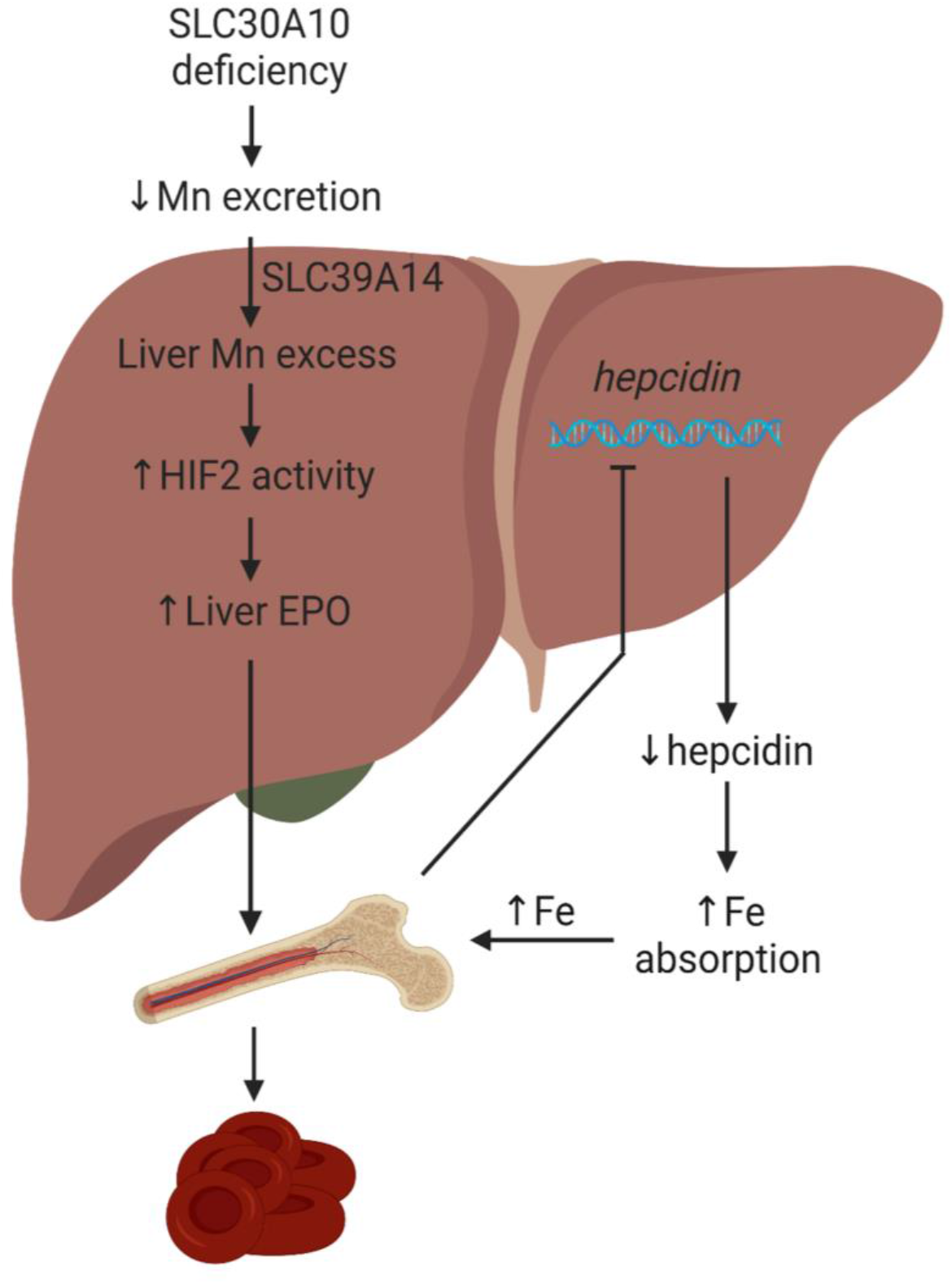

## Introduction

Manganese (Mn) is an essential yet potentially toxic metal (1, 2). It is an essential enzymatic cofactor in multiple biological processes including antioxidant defense and protein glycosylation. When present in excess, Mn is toxic, with toxicity attributed to factors such as oxidative stress and mitochondrial dysfunction. Mn toxicity is typically caused by environmental exposure and manifests as neurologic impairment. Inherited causes of Mn excess are rare, but study of these diseases has greatly informed our understanding of the molecular basis of Mn homeostasis in the human body.

Mutations in metal transport proteins SLC30A10 (ZNT10) and SLC39A14 (ZIP14) are two recently described inherited causes of Mn excess. Patients with SLC30A10 deficiency were first reported in 2012, while patients with SLC39A14 deficiency were first reported in 2016 (3). SLC30A10 is a apical membrane protein that exports Mn from hepatocytes into bile and from enterocytes into the lumen of the gastrointestinal tract (4–6). SLC30A10 deficiency impairs gastrointestinal Mn excretion, resulting in severe Mn excess as well as neurologic and liver dysfunction, polycythemia, and erythropoietin (EPO) excess. Neurologic and liver disease are attributed to Mn toxicity. Polycythemia is attributed to EPO excess but the basis of EPO excess in this disease has yet to be established. SLC39A14 is a basolateral membrane protein that imports Mn from liver sinusoids into hepatocytes and from blood into enterocytes (7–13). Like SLC30A10 deficiency, SLC39A14 deficiency impairs gastrointestinal excretion, resulting in severe Mn excess and neurologic deficits. Unlike SLC30A10 deficiency, SLC39A14 deficiency does not produce liver disease, polycythemia, or EPO excess.

As stated above, the basis of EPO excess in SLC30A10 deficiency is unknown. *EPO* is the canonical hypoxia-regulated gene. Under conditions of hypoxia, the kidneys secrete EPO to stimulate erythropoiesis in a process dependent upon transcription factors known as hypoxia-inducible factors (HIFs) (14). HIFs consist of labile α subunits and constitutively expressed β subunits. Under conditions of normoxia, HIF α subunits are subjected to prolyl hydroxylation by prolyl hydroxylases (PHDs), followed by ubiquitination and degradation. Under conditions of hypoxia, HIF α subunits are spared from degradation and can activate transcription of *EPO* and a variety of other genes. While a role for HIFs in EPO excess and polycythemia in SLC30A10 deficiency has yet to be explored, several studies have shown links between Mn, EPO, and HIFs. Mn treatment of human airway cell line Hep2 increases HIF1α protein levels by activating MAPKs (15), although the validity of this cell line has been called into question recently (16). Mn treatment of lung cancer cells inhibits PHD activity, increases HIF1α protein levels, and stimulates expression of vascular endothelial growth factor (VEGF), a known HIF target gene (17). Mn treatment of human pulmonary epithelial cell lines increases VEGF expression, and Mn inhalation in mice increases pulmonary expression of VEGF and multiple other angiogenesis-associated genes (18). Mn treatment of human hepatoma cell line Hep3B stimulates *EPO* expression (19). A recent link between Mn, SLC30A10, and HIFs has been reported as well—Mn treatment of hepatic cell lines increases HIF1- and HIF2-dependent SLC30A10 expression by inhibiting prolyl hydroxylation of HIFs (20).

In this study, we investigate the link between SLC30A10 deficiency and EPO excess using Slc30a10-deficient mouse models and dietary, genetic, pharmacologic, and transcriptomic approaches. As described below, we demonstrate that hepatic Hif1 has minimal impact on Epo excess and polycythemia, while hepatic Hif2 not only drives Epo excess and polycythemia but also contributes to aberrant hepatic gene expression, increased dietary iron absorption, and systemic Mn excess in Slc30a10-deficient mice. We also discuss implications of our observations for our understanding and treatment of SLC30A10 deficiency.

## Results

### Slc30a10^-/-^ mice develop Mn-dependent Epo excess and polycythemia

To investigate the basis of EPO excess in SLC30A10 deficiency, we first measured Epo levels in our previously described *Slc30a10*^*-/-*^ mouse model (6). Serum Epo levels were increased in female and male mutant mice (Fig. 1A). To identify a potential source of excess Epo, *Epo* RNA levels were assessed and found to be decreased in kidneys yet increased in livers of *Slc30a10*^*-/-*^ mice (Fig. 1B). This suggested that the liver is the source of excess EPO in SLC30A10 deficiency. (We attempted to measure *EPO* RNA levels in liver biopsies from patients with SLC30A10 mutations (21, 22), but liver biopsies yielded inadequate quality RNA for analysis.) Given that EPO is a hypoxia-regulated gene, we next used PCR arrays to assess expression of other hypoxia-regulated genes. 24% of 93 hypoxia-regulated genes, including *Epo, Serpine1* (serpin family E member 1), *Hk2* (hexokinase 2), and *Anxa2* (annexin A2), were upregulated at least two-fold in mutant livers (Fig. 1C, D). Upregulation of *Serpine1, Hk2*, and *Anxa2*, the top three upregulated genes after *Epo*, was validated by qPCR in *Slc30a10*^*-/-*^ livers (Fig. 1E). We next determined if metal excess contributed to Epo excess in mutant mice. Excess cobalt can stimulate hypoxia-regulated gene expression in the absence of hypoxia (23). We measured liver cobalt levels in *Slc30a10*^*-/-*^ mice, but levels were decreased in female mice and unchanged in male mice (Fig. 1F). This suggested that cobalt excess was not driving Epo excess in *Slc30a10*^*-/-*^ mice. To determine if Mn excess contributed to Epo excess and polycythemia in mutant mice, we weaned *Slc30a10*^*-/-*^ mice onto Mn-deficient diets and measured liver Mn levels, liver *Epo* RNA levels, and red blood cell (RBC) counts in six-week-old mice. Liver Mn and *Epo* RNA levels and RBC counts were decreased in mutant mice weaned onto Mn-deficient diets (Fig. 1G-I). This indicated that Mn excess plays a causal role in Epo excess and polycythemia in *Slc30a10*^*-/-*^ mice.

**Fig. 1:**
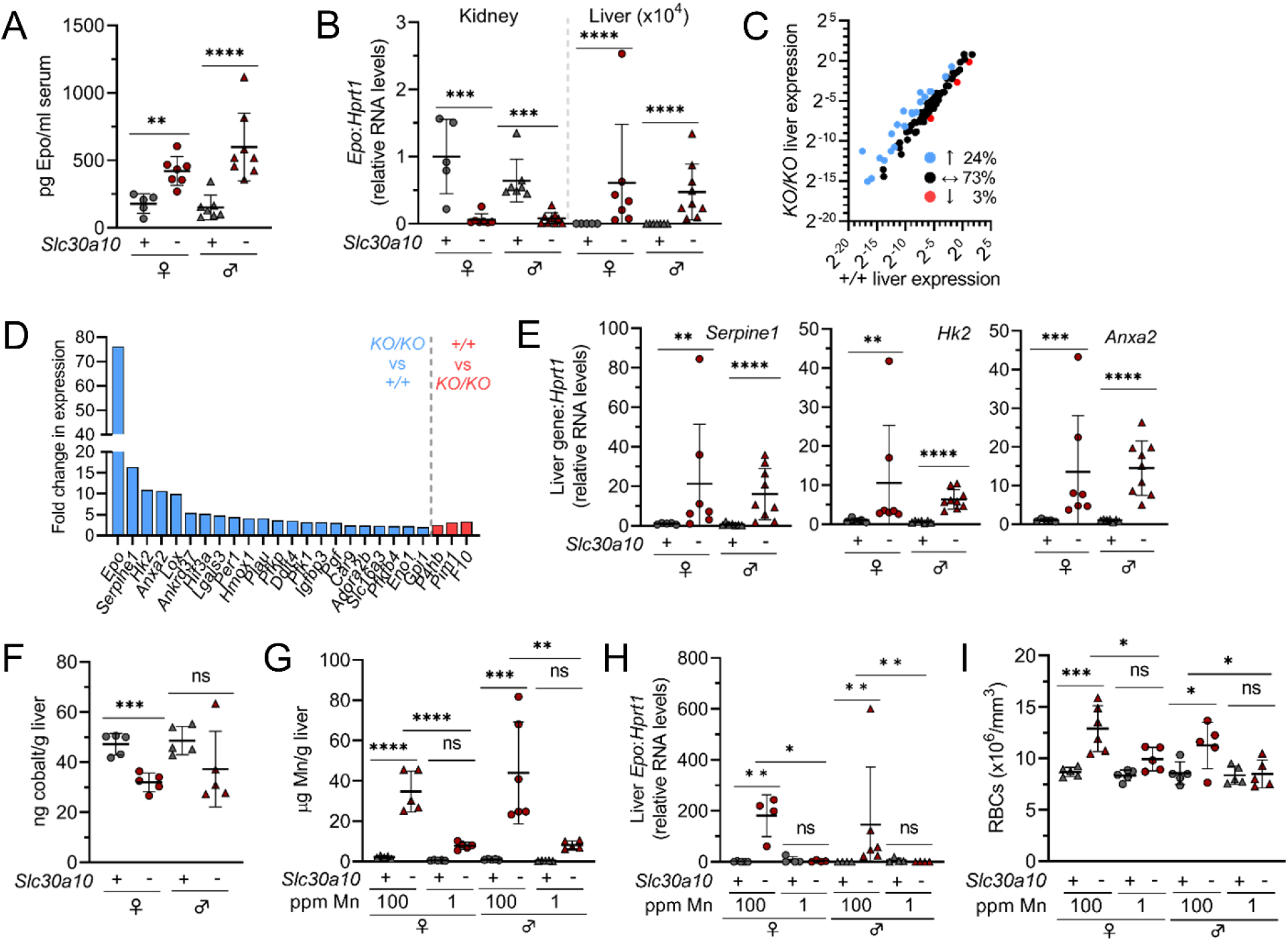
*Slc30a10*^*-/-*^ mice develop Mn-dependent Epo excess and polycythemia. **(A, B)** Two-month-old *Slc30a10*^*+/+*^ and *Slc30a10*^*-/-*^ mice were analyzed for serum Epo levels by ELISA (**A**) and kidney and liver *Epo* RNA levels by qPCR (**B**). **(C-E)** Livers from two-month-old *Slc30a10*^*+/+*^ and *Slc30a10*^*-/-*^ mice were analyzed by PCR array for hypoxia-regulated genes up- (blue) or down- (red) regulated at least two-fold (**C, D**), followed by qPCR validation of top three differentially regulated genes after *Epo* (**E**). **(F)** Two-month-old *Slc30a10*^*+/+*^ and *Slc30a10*^*-/-*^ mice were analyzed for liver cobalt levels by GFAAS. **(G-I)** *Slc30a10*^*+/+*^ and *Slc30a10*^*-/-*^ mice were weaned onto Mn-sufficient (100 ppm) or –deficient (1 ppm) diets, then analyzed at six weeks for liver Mn levels by ICPES (**G**), RBC counts by complete blood counts (**H**), and liver *Epo* RNA levels by qPCR (**I**). Data are represented as means +/- standard deviation, with at least four animals per group, except for (**C, D**) where three mice were used. In this figure and all other figures, data were tested for normal distribution by Shapiro-Wilk test; if not normally distributed, data were log transformed. Two groups were compared by unpaired, two-tailed t test. Three or more groups (within each sex) were compared by one-way ANOVA with Tukey’s multiple comparisons test. (ns P=>0.05, * P<0.05, ** P<0.01, *** P<0.001, **** P<0.0001)

### Slc39a14 deficiency corrects liver Epo excess and polycythemia in Slc30a10^-/-^ mice

To further investigate the basis of EPO excess and polycythemia in SLC30A10 deficiency, we interrogated SLC39A14. Essential for gastrointestinal Mn excretion, SLC39A14 imports Mn from blood into hepatocytes and enterocytes. Both SLC30A10 and SLC39A14 deficiency cause Mn excess and neurologic disease but SLC39A14 deficiency is not associated with liver disease, liver Mn excess, or polycythemia. Given the absence of liver Mn excess and polycythemia in SLC39A14 deficiency, we determined the impact of Slc39a14 deficiency on *Slc30a10*^*-/-*^ mice by generating and characterizing Slc30a10- and Slc39a14-deficient mice (*Slc30a10*^*-/-*^ *Slc39a14*^*-/-*^) (Fig. 2A, B). Slc39a14 deficiency normalized liver Mn levels, indicating that Slc39a14 is essential for Mn excess in Slc30a10 deficiency as previously shown (24) (Fig. 2C). Slc39a14 deficiency also normalized liver *Epo, Serpine1, Hk2*, and *Anxa2* RNA levels, RBC counts, hemoglobin levels, and hematocrits in *Slc30a10*^*-/-*^ mice (Fig. 2D-J). This indicated that Slc39a14 is also essential for aberrant expression of several hypoxia-regulated genes and polycythemia in Slc30a10 deficiency. We next assessed *Epo* RNA and Mn levels in kidneys. In contrast to its impact on liver *Epo* RNA and Mn levels, Slc39a14 deficiency had no impact on kidney *Epo* RNA levels and increased kidney Mn levels in *Slc30a10*^*-/-*^ mice (Fig. 2K, L). Finally, we measured Mn levels in other extrahepatic sites/tissues. Slc39a14 deficiency did not change blood or pancreas Mn levels but did increase bone, brain, and heart Mn levels and decrease small and large intestine Mn levels in *Slc30a10*^*-/-*^ mice (Fig. 2M-S). Overall, characterization of mice with Slc39a14 and Slc30a10 deficiency indicated that Slc39a14 is essential for development of liver Mn excess, *Epo* excess, and polycythemia in *Slc30a10*^*-/-*^ mice. This suggested that SLC39A14-dependent import of Mn into liver drives EPO excess and polycythemia in SLC30A10 deficiency. However, given that Slc39a14 deficiency also impacted Mn levels in extrahepatic tissues, we cannot formally exclude a role for SLC39A14 in those other tissues in determining EPO excess and polycythemia.

**Fig. 2:**
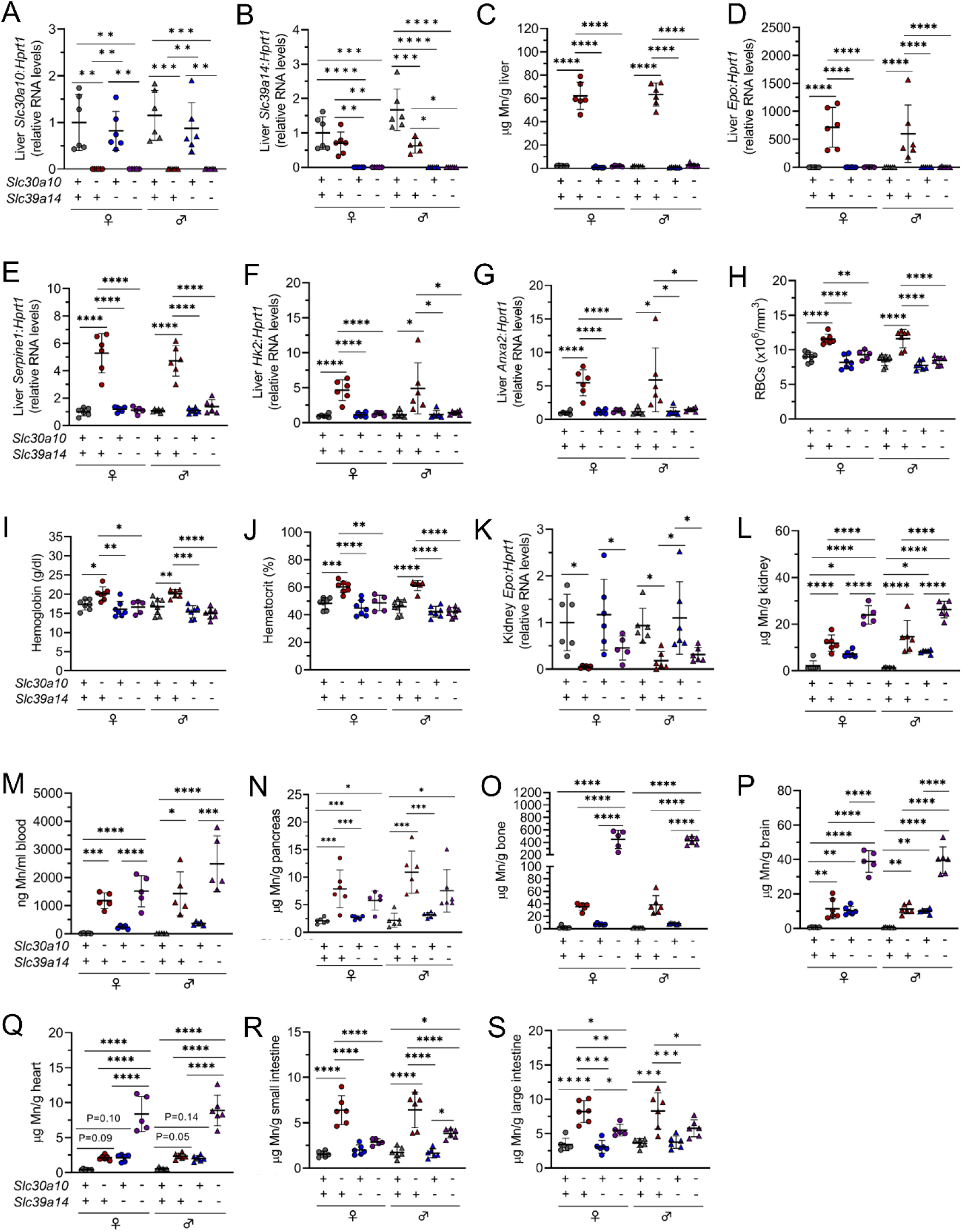
Slc39a14 deficiency corrects liver *Epo* excess and polycythemia in *Slc30a10*^*-/-*^ mice. Five-week-old *Slc30a10 Slc39a14* mice were analyzed for: liver *Slc30a10* (**A**) and *Slc39a14* (**B**) RNA levels by qPCR; liver Mn levels by ICPES (**C**); liver *Epo* (**D**), *Serpine1* (**E**), *Hk2* (**F**), and *Anxa2* (**G**) RNA levels by qPCR; RBC counts (**H**), hemoglobin levels (**I**), and hematocrits (**J**) by complete blood counts; kidney *Epo* RNA levels by qPCR (**K**); kidney Mn levels by ICPES (**L**); blood Mn levels by GFAAS (**M**); pancreas (**N**), bone (**O**), brain (**P**), heart (**Q**), small intestine (**R**), and large intestine (**S**) Mn levels by ICPES. Data are represented and statistics performed and annotated as in Fig. 1.

### Hif2a antisense oligonucleotides (ASOs) decrease liver Epo RNA levels in Slc30a10^-/-^ mice

*EPO* is the canonical hypoxia-regulated gene, and HIFs are essential mediators of the cellular response to hypoxia. To determine if Hifs mediate Epo excess in *Slc30a10*^*-/-*^ mice, we employed two approaches. In the first, we treated three-week-old mutant mice with GalNac-conjugated control, *Hif1a*, or *Hif2a* ASOs for three weeks. We measured liver *Slc30a10* RNA levels and observed no impact of ASO treatment on *Slc30a10* RNA levels (Fig. 3A). We next assessed liver *Hif* RNA levels. Surprisingly, liver *Hif1a* and *Hif2a* RNA levels were decreased in untreated *Slc30a10*^*-/-*^ mice (Fig. 3B, C). *Hif1a* ASOs decreased liver *Hif1a* and *Hif2a* RNA levels beyond the decrease observed in untreated *Slc30a10*^*-/-*^ mice (Fig. 3B, C). In contrast, *Hif2a* ASOs increased liver *Hif1a* RNA levels, while they had no impact on liver *Hif2a* RNA levels possibly because *Hif2a* RNA levels were already low in untreated *Slc30a10*^*-/-*^ mice (Fig. 3B, C). Measurement of kidney *Hif1a* and *Hif2a* RNA levels revealed no changes in ASO-treated mice (Fig. 3D, E). We next assessed the impact of ASOs on *Epo* levels and RBC parameters. *Hif2a* ASOs, but not *Hif1a* ASOs, decreased liver *Epo* RNA levels, while neither ASO impacted kidney *Epo* RNA levels or RBC parameters (Fig. 3F-J). (The decrease in liver *Epo* RNA levels in *Hif1a* ASO-treated mice was not significant.) We also observed that *Hif2a* ASOs increased body mass in *Slc30a10*^*-/-*^ mice (Fig. 3K). Finally, we noted that *Hif1a* ASOs decreased bile flow rates while neither ASO altered bile or liver Mn levels (Fig. 3K-N). (Despite severe Mn excess, bile Mn levels are not elevated in *Slc30a10*^*-/-*^ mice, which we attribute to Slc30a10’s essential role in hepatobiliary Mn excretion.) Taken together, these data suggested that Hif2, but not Hif1, contributes to Epo excess in *Slc30a10*^*-/-*^ mice.

**Fig. 3:**
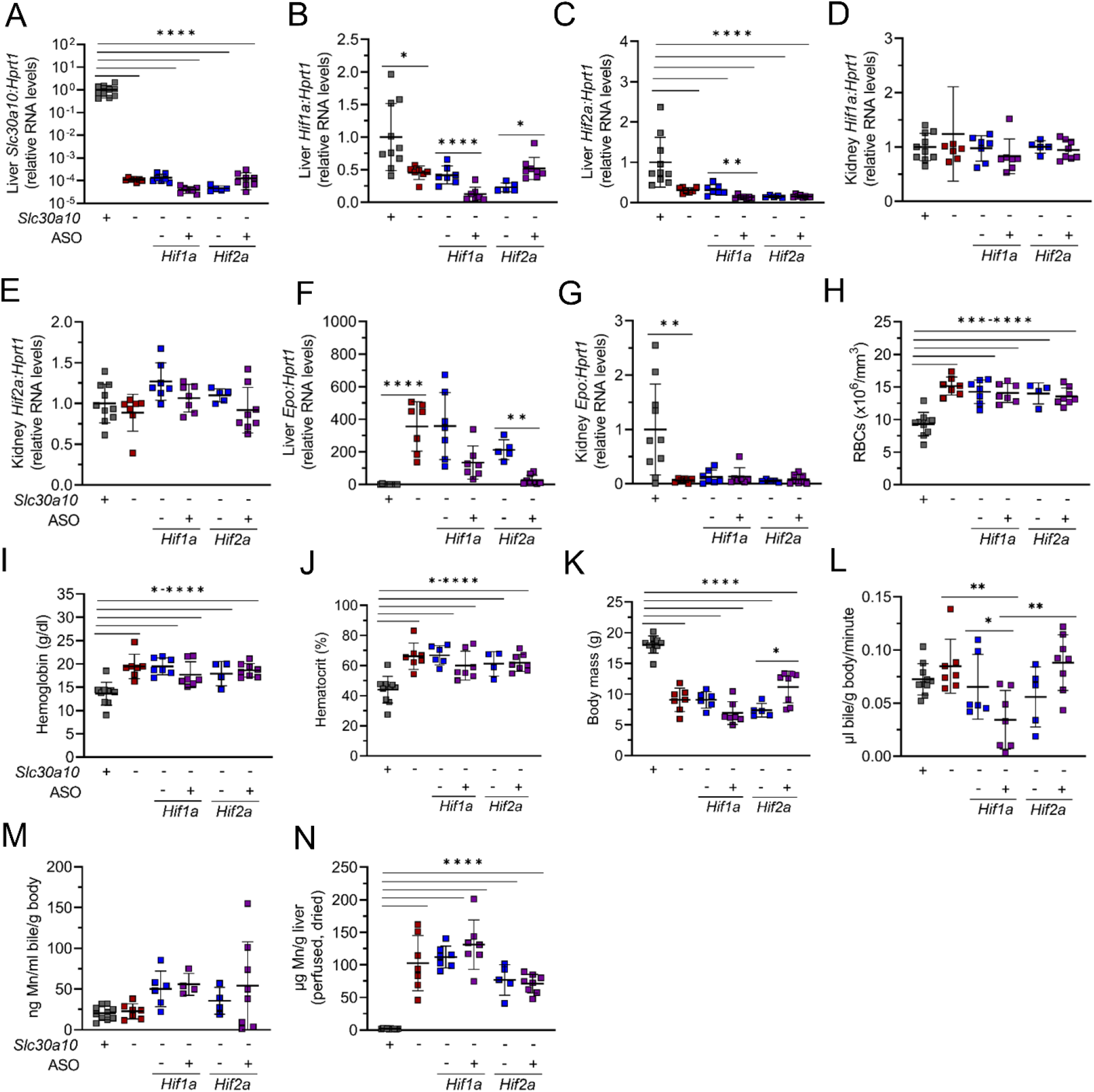
*Hif2a* ASOs decrease liver *Epo* RNA levels in *Slc30a10*^*-/-*^ mice. Weanling *Slc30a10*^*+/+*^ mice were treated with saline and *Slc30a10*^*-/-*^ mice with GalNAc-conjugated control, *Hif1a*, or *Hif2a* ASOs twice a week for three weeks. Mice were then analyzed for: liver *Slc30a10* RNA levels (**A**), liver *Hif1a* (**B**) and *Hif2a* (**C**) RNA levels, kidney *Hif1a* (**D**) and *Hif2a* (**E**) RNA levels, and liver (**F**) and kidney (**G**) *Epo* RNA levels by qPCR; RBC counts (**H**), hemoglobin levels (**I**), and hematocrits (**J**) by complete blood counts; body mass (**K**); bile flow rates (**L**); bile Mn levels by GFAAS (**M**); and liver Mn levels by ICPES (**N**). Data are represented and statistics performed and annotated as in Fig. 1.

### Hepatocyte deficiency in Hif2a but not Hif1a corrects Epo excess and polycythemia and decreases Mn levels in Slc30a10^-/-^ mice

In our second approach to determine if Hifs mediate Epo excess in *Slc30a10*^*-/-*^ mice, we generated two-month-old *Slc30a10*^*-/-*^ mice with hepatocyte *Hif1a* or *Hif2a* deficiency using an albumin promoter-driven Cre recombinase transgene (*Alb*). We first characterized *Slc30a10 Hif1a Alb* mice. Hepatocyte Hif1a deficiency had no impact on liver *Slc30a10* RNA levels (Fig. 4A). As observed above, liver *Hif1a* and *Hif2a* RNA levels were decreased in *Slc30a10*^*-/-*^ mice (Fig. 4B, C). Hepatocyte Hif1a deficiency decreased liver *Hif1a* RNA levels but had no effect on liver *Hif2a* RNA levels or kidney *Hif1a* or *Hif2a* RNA levels, confirming the specificity of the *Hif1a*^*fl/fl*^ *Alb* approach (Fig. 4B-E). Hepatocyte Hif1a deficiency also did not impact the increased liver *Epo* RNA levels, decreased kidney *Epo* RNA levels, decreased body mass (in male mice), or aberrant RBC parameters in *Slc30a10*^*-/-*^ mice (Fig. 4F-K). Hepatocyte Hif1a deficiency also had no impact on bile flow rates or bile, liver, or blood Mn levels in *Slc30a10*^*-/-*^ mice, although bile flow rates were increased in male *Slc30a10*^*+/+*^ *Hif1a*^*fl/fl*^ *Alb* mice and liver Mn levels were increased in male *Slc30a10*^*-/-*^ *Hif1a*^*fl/fl*^ *Alb* mice (Fig. 4L-O). Overall, our characterization of *Slc30a10 Hif1a Alb* mice indicated that hepatocyte Hif1a plays a minimal role in phenotypes assessed in *Slc30a10*^*-/-*^ mice.

**Fig. 4:**
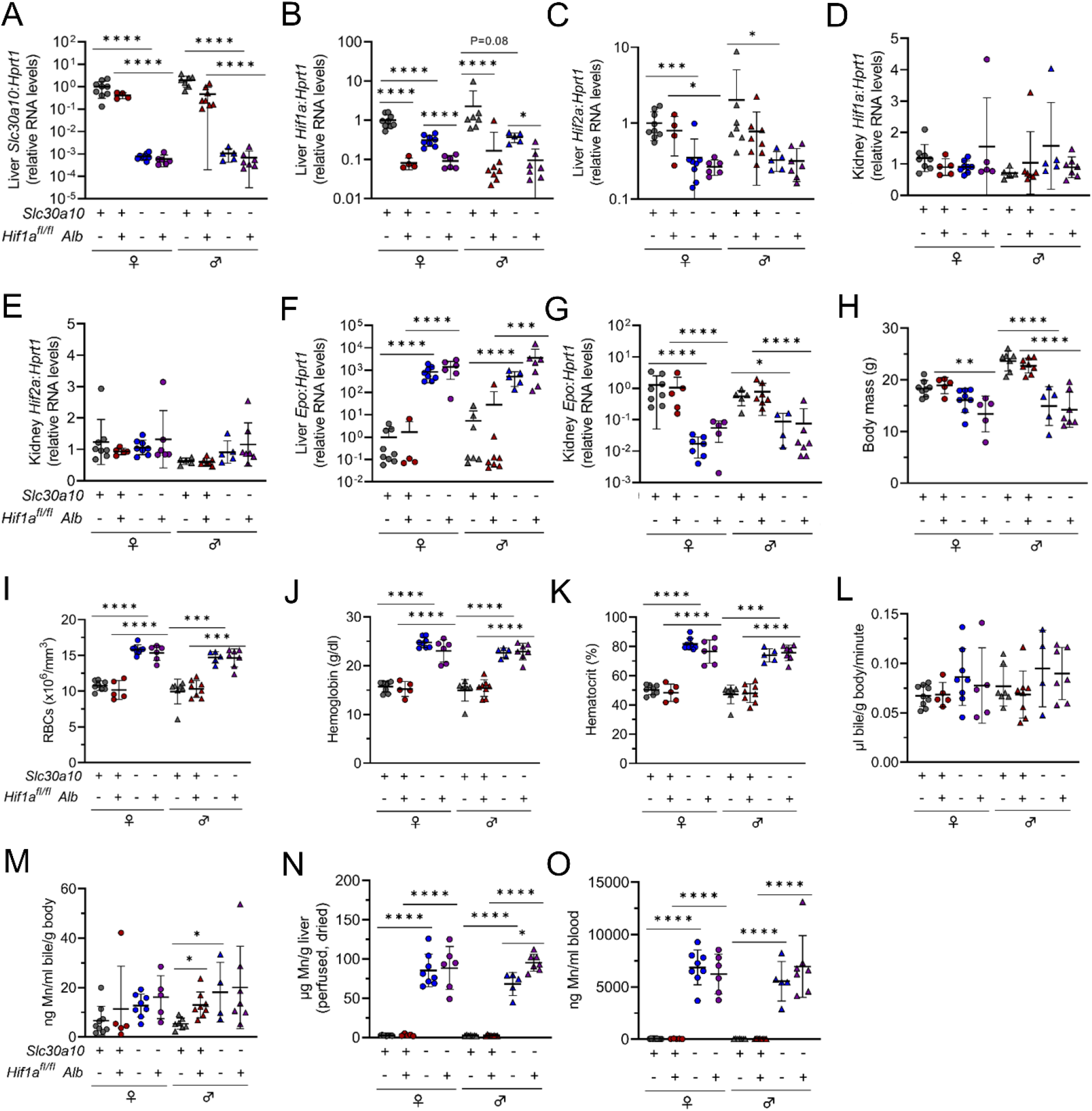
Hepatocyte Hif1a deficiency does not impact *Epo* levels or polycythemia in *Slc30a10*^*-/-*^ mice. Two-month-old *Slc30a10 Hif1a*^*fl/fl*^ *+/-Alb* mice were analyzed for: liver *Slc30a10* (**A**), *Hif1a* (**B**), and *Hif2a* (**C**) RNA levels, kidney *Hif1a* (**D**) and *Hif2a* (**E**) RNA levels, and liver (**F**) and kidney (**G**) *Epo* RNA levels by qPCR; body mass (**H**); RBC counts (**I**), hemoglobin levels (**J**), and hematocrits (**K**) by complete blood counts; bile flow rates (**L**); bile Mn levels by GFAAS (**M**); liver Mn levels by ICPES (**N**); blood Mn levels by GFAAS (**O**). Data are represented and statistics performed and annotated as in Fig. 1.

We next characterized *Slc30a10 Hif2a Alb* mice. Hepatocyte Hif2a deficiency had no impact on liver *Slc30a10* RNA levels (Fig. 5A). Liver *Hif1a* and *Hif2a* RNA levels were decreased in *Slc30a10*^*-/-*^ mice as noted earlier (Fig. 5B, C). Hepatocyte Hif2a deficiency decreased liver *Hif2a* but not *Hif1a* RNA levels (Fig. 5B, C). Hepatocyte Hif2a deficiency had no impact on kidney *Hif2a* or *Hif1a* RNA levels except for decreased kidney *Hif2* RNA levels in male *Slc30a10*^*-/-*^ mice (Fig. 5D, E). We did observe that kidney *Hif2a* RNA levels were increased in *Slc30a10*^*-/-*^ *Hif2a*^*fl/fl*^ mice relative to *Slc30a10*^*+/+*^ *Hif2a*^*fl/fl*^ mice, but this was not observed in *Slc30a10* cohorts analyzed in Figs. 2 and 3 (Fig. 5D). In contrast to hepatocyte Hif1a deficiency, hepatocyte Hif2a deficiency decreased liver *Epo* RNA levels and increased kidney *Epo* RNA levels in *Slc30a10*^*-/-*^ mice (Fig. 5F, G). Hepatocyte Hif2a deficiency also increased body mass in female *Slc30a10*^*-/-*^ mice (and insignificantly in male mutant mice) and normalized RBC parameters in all *Slc30a10*^*-/-*^ mice (Fig. 5H-K). We next assessed bile, blood, and tissue Mn levels. Hepatocyte Hif2a deficiency had no impact on bile flow rates but did decrease Mn levels in bile, liver, bone, brain, pancreas (in females), kidney (in females), and blood in *Slc30a10*^*-/-*^ mice, although decreases in blood were not significant (Fig. 5L-S). Taken together, these data indicated that Hif2a is not only a key determinant of Epo excess and polycythemia but also Mn excess in *Slc30a10*^*-/-*^ mice.

**Fig. 5:**
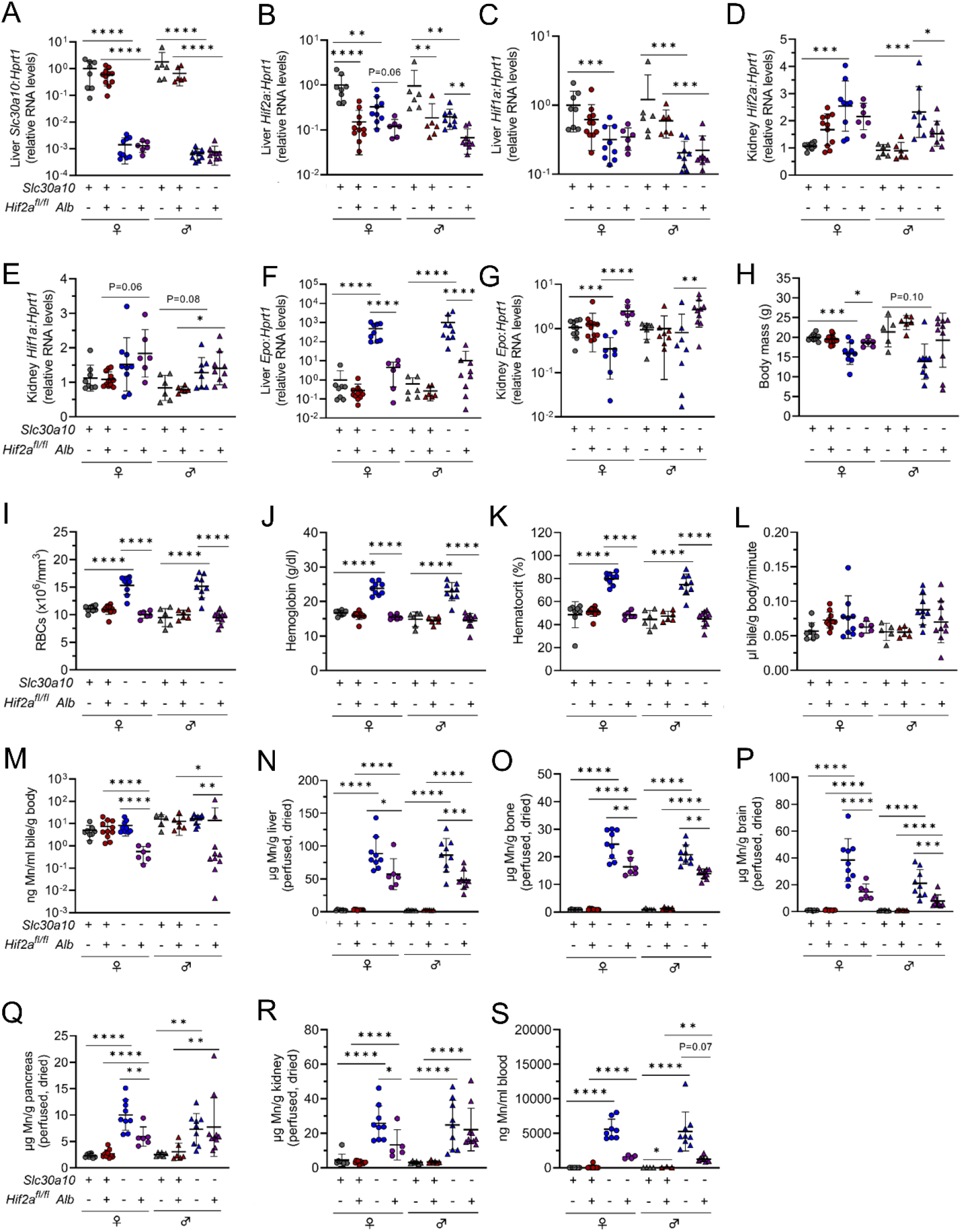
Hepatocyte Hif2a deficiency corrects *Epo* excess and polycythemia and decreases Mn levels in *Slc30a10*^*-/-*^ mice. Two-month-old *Slc30a10 Hif2a*^*fl/fl*^ +/-*Alb* mice were analyzed for: liver *Slc30a10* (**A**), *Hif2a* (**B**), and *Hif1a* (**C**) RNA levels, kidney *Hif2a* (**D**) and *Hif1a* (**E**) RNA levels, and liver (**F**) and kidney (**G**) *Epo* RNA levels by qPCR; body mass (**H**); RBC counts (**I**), hemoglobin levels (**J**), and hematocrits (**K**) by complete blood counts; bile flow rates (**L**); bile Mn levels by GFAAS (**M**); liver (**N**), bone (**O**), brain (**P**), pancreas (**Q**), and kidney (**R**) Mn levels by ICPES; blood Mn levels by GFAAS (**S**). Data are represented and statistics performed and annotated as in Fig. 1.

We next considered the extent to which correction of polycythemia by hepatic Hif2a deficiency in *Slc30a10*^*-/-*^ mice contributed to decreased tissue Mn levels shown above. To clear blood from tissues so that tissue Mn levels would reflect parenchymal but not blood Mn, *Slc30a10 Hif1a* and *Slc30a10 Hif2a* mice were perfused with saline prior to tissue harvest, and tissues were dried before metal measurements. To assess the efficacy of perfusion on clearing blood from tissues, we focused on tissue iron levels. We previously demonstrated that *Slc30a10*^*-/-*^ mice (without perfusion prior to tissue harvest) have increased total iron levels in liver (data regraphed for reference in Fig. S1A) (6). However, liver non-heme iron levels did not differ between *Slc30a10*^*+/+*^ and *Slc30a10*^*-/-*^ mice (Fig. S1B). This indicated that the increased total iron levels reflected RBC hemoglobin, not parenchymal iron, in the liver. We next analyzed total and non-heme iron levels in livers from perfused *Slc30a10 Hif1a* and *Slc30a10 Hif2a* mice. Despite the presence of polycythemia in *Slc30a10*^*-/-*^ *Hif1a*^*fl/fl*^ and *Slc30a10*^*-/-*^ *Hif1a*^*fl/fl*^ *Alb* mice, neither total nor non-heme iron levels were increased in livers from these mice relative to their *Slc30a10*^*+/+*^ counterparts (Fig. S1C, D). Similarly, despite the presence of polycythemia in *Slc30a10*^*-/-*^ *Hif2a*^*fl/fl*^ mice, neither total nor non-heme iron levels were increased in livers from these mice relative to *Slc30a10*^*+/+*^ *Hif2a*^*fl/fl*^ mice (Fig. S1E, F). These findings suggested that perfusion was effective at clearing blood from livers in polycythemic mice. (Non-heme iron levels were decreased in *Slc30a10*^*-/-*^ *Hif1a*^*fl/fl*^ *Alb* livers relative to *Slc30a10*^*-/-*^ *Hif1a*^*fl/fl*^ livers, while total and non-heme iron levels were increased in *Slc30a10*^*-/-*^ *Hif2a*^*fl/fl*^ *Alb* livers relative to *Slc30a10*^*+/+*^ *Hif2a*^*fl/fl*^ *Alb* livers in female mice—the relevance of this is not clear at this time.) In our previously published study, we also observed that total iron levels were increased in pancreas, kidney, and brain in *Slc30a10*^*-/-*^ mice (data regraphed for reference in Fig. S1G-I) (6). Total iron levels in pancreas, kidney, or brain (males only) did not differ by genotype for *Slc30a10 Hif2a* mice, suggesting that perfusion was effective at clearing blood from these tissues (Fig. S1J-L). However, total iron levels were increased in *Slc30a10*^*-/-*^ *Hif2a*^*fl/fl*^ brains relative to *Slc30a10*^*+/+*^ *Hif2a*^*fl/fl*^ brains for female mice (Fig. S1L).

### Slc30a10 deficiency prominently impacts hepatic gene expression

To further interrogate the role of hepatic Hifs in Slc30a10 deficiency, we next identified Hif-dependent genes aberrantly expressed in *Slc30a10*^*-/-*^ mice. To do this, we performed bulk RNA-seq on livers from *Slc30a10 Hif1a* and *Slc30a10 Hif2a* mice analyzed in Figs. 4 and 5. To determine if we could pool *Hif1a*^*fl/fl*^ and *Hif2a*^*fl/fl*^ genotypes in this analysis, we compared gene expression between these two groups and found that fewer than 25 of more than 13,000 genes detected were differentially expressed between *Hif1a*^*fl/fl*^ and *Hif2a*^*fl/fl*^ mice on *Slc30a10*^*+/+*^ or *Slc30a10*^*-/-*^ backgrounds (Fig. S2). This indicated that the floxed alleles of *Hif1a* and *Hif2a* in absence of a Cre recombinase transgene had minimal impact on hepatic gene expression. We next established the impact of Slc30a10 deficiency on hepatic gene expression by comparing *Slc30a10*^*-/-*^ and *Slc30a10*^*+/+*^ livers, with *Hif1a*^*fl/fl*^ and *Hif2a*^*fl/fl*^ samples pooled. Samples grouped largely by *Slc30a10* genotype (Fig. 6A-D). 1334 genes in female mice and 2246 genes in male mice were differentially expressed in *Slc30a10*^*-/-*^ versus *Slc30a10*^*+/+*^ livers (Fig. 6E, F). Included in these differentially expressed genes were *Slc30a10* (downregulated as expected in *Slc30a10*^*-/-*^ mice), *Serpine1, Hk2, Anxa2* (upregulated, as shown earlier by PCR array in Fig. 1C-E), and hepcidin (*Hamp*; downregulated; discussed below), but not *Epo* (also discussed below). Of all the genes differentially expressed in female and male mice, 880 genes were differentially expressed in both sexes and aligned largely with cell cycle (Fig. S3). 454 genes were differentially expressed only in females and aligned mainly with DNA replication/cell cycle (Fig. S4). Most of the cell cycle-aligned genes were upregulated in *Slc30a10*^*-/-*^ livers (Fig. S3D, S4D). 1365 genes were differentially expressed only in males and aligned mostly with cell motility (Fig. S5).

**Fig. 6:**
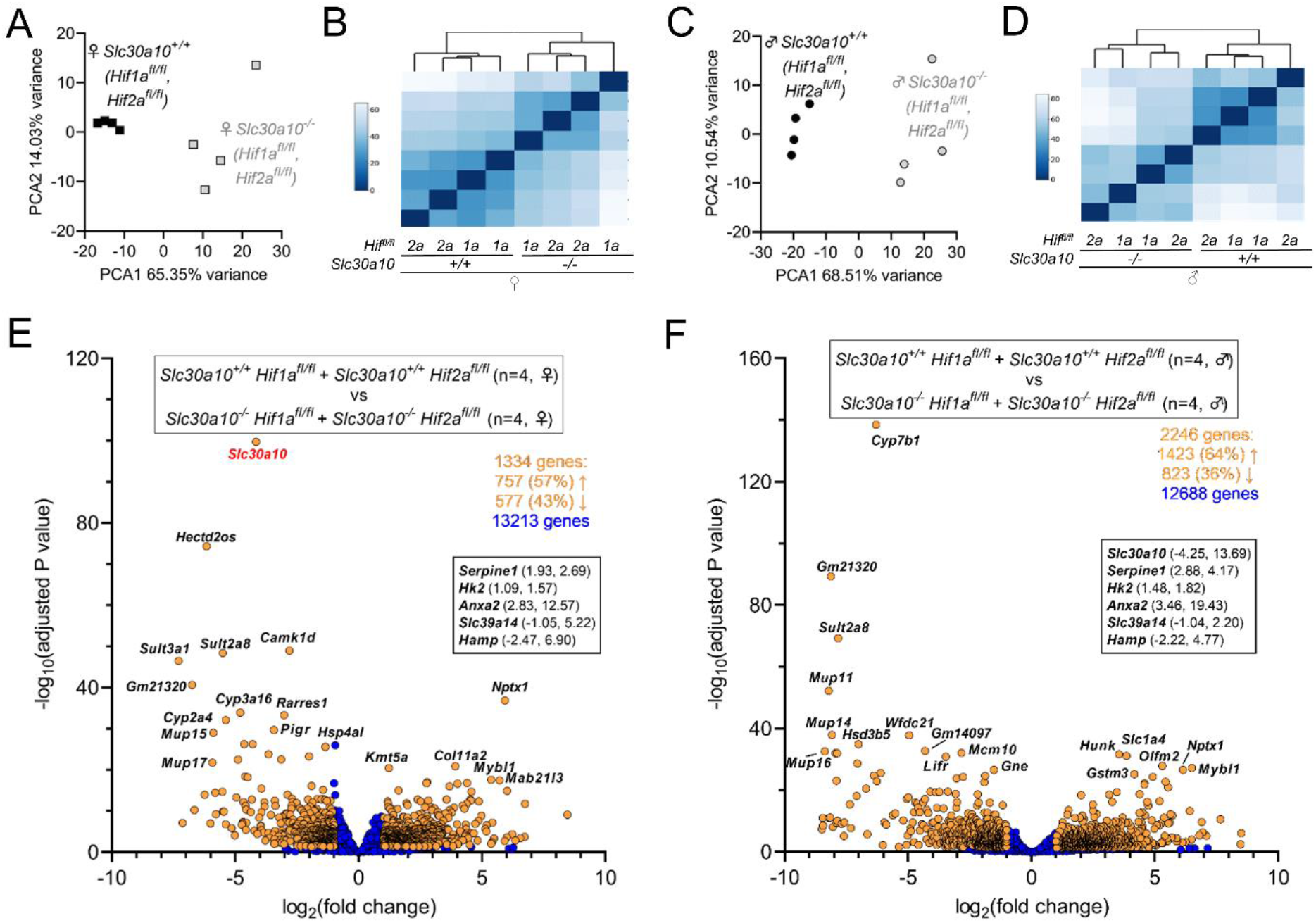
Slc30a10 deficiency prominently impacts hepatic gene expression. Principal component (**A, B**) and similarity analysis (**C, D**) of samples and volcano plots (**E, F**) of genes differentially expressed between *Slc30a10*^*+/+*^ *Hif1a*^*fl/fl*^ or *Hif2a*^*fl/fl*^ mice and *Slc30a10*^*-/-*^ *Hif1a*^*fl/fl*^ or *Hif2a*^*fl/fl*^ mice; female (**A, B, E**) and male (**C, D, F**). In volcano plots, differentially expressed genes (adjusted P value<0.05 and absolute value of log_2_(fold change)>1) are shown as light orange points with gene names shown adjacent as space permitted; non-differentially expressed genes are shown as blue points; x-y coordinates of additional genes of interest shown in smaller box. Genes with log_2_(fold change)<0 are more abundantly expressed in first group listed in box at top of plot; genes with log_2_(fold change)>0 are more abundantly expressed in second group listed.

### Hepatic deficiency in Hif2a but not Hif1a has a prominent impact on hepatic gene expression in Slc30a10^-/-^ mice

After identifying liver genes differentially expressed in Slc30a10 deficiency, we next determined which of those genes were differentially expressed in a Hif-dependent manner. We first focused on Hif1a by performing two comparisons: *Slc30a10*^*-/-*^ *Hif1a*^*fl/fl*^ versus *Slc30a10*^*+/+*^ *Hif1a*^*fl/fl*^ livers to identify genes differentially expressed in Slc30a10 deficiency in the *Slc30a10 Hif1a* cohort, and *Slc30a10*^*-/-*^ *Hif1a*^*fl/fl*^ *Alb* versus *Slc30a10*^*-/-*^ *Hif1a*^*fl/fl*^ livers to identify genes differentially expressed in a Hif1a-dependent manner in Slc30a10 deficiency. For the *Slc30a10*^*-/-*^ *Hif1a*^*fl/fl*^ versus *Slc30a10*^*+/+*^ *Hif1a*^*fl/fl*^ comparison, female and male *Slc30a10*^*-/-*^ *Hif1a*^*fl/fl*^ samples grouped together, while *Slc30a10*^*+/+*^ *Hif1a*^*fl/fl*^ samples grouped by sex but still separately from *Slc30a10*^*-/-*^ *Hif1a*^*fl/fl*^ samples (Fig. 7A, B). In contrast, for the *Slc30a10*^*-/-*^ *Hif1a*^*fl/fl*^ *Alb* versus *Slc30a10*^*-/-*^ *Hif1a*^*fl/fl*^ comparison, samples did not group by genotype or sex (Fig. 7C, D). 1528 genes, including *Slc30a10* (downregulated in *Slc30a10*^*-/-*^ livers), *Hk2, Anxa2* (both upregulated), and *Hamp* (downregulated), were differentially expressed between *Slc30a10*^*-/-*^ *Hif1a*^*fl/fl*^ and *Slc30a10*^*+/+*^ *Hif1a*^*fl/fl*^ livers (Fig. 7E). These genes aligned largely with metabolic processes and responses to stimuli, with most of metabolic process-aligned genes downregulated in *Slc30a10*^*-/-*^ livers and most of the stimuli response-aligned genes upregulated in *Slc30a10*^*-/-*^ livers (Fig. S6). In contrast to the *Slc30a10*^*-/-*^ *Hif1a*^*fl/fl*^ *Alb* versus *Slc30a10*^*+/+*^ *Hif1a*^*fl/fl*^ analysis, only 21 genes were differentially expressed between *Slc30a10*^*-/-*^ *Hif1a*^*fl/fl*^ *Alb* and *Slc30a10*^*-/-*^ *Hif1a*^*fl/fl*^ livers (Fig. 7F). The minimal impact of hepatic Hif1a deficiency on differential gene expression in *Slc30a10*^*-/-*^ livers was consistent with the minimal impact of hepatic Hif1a deficiency on phenotypes assessed in Fig. 4.

**Fig. 7:**
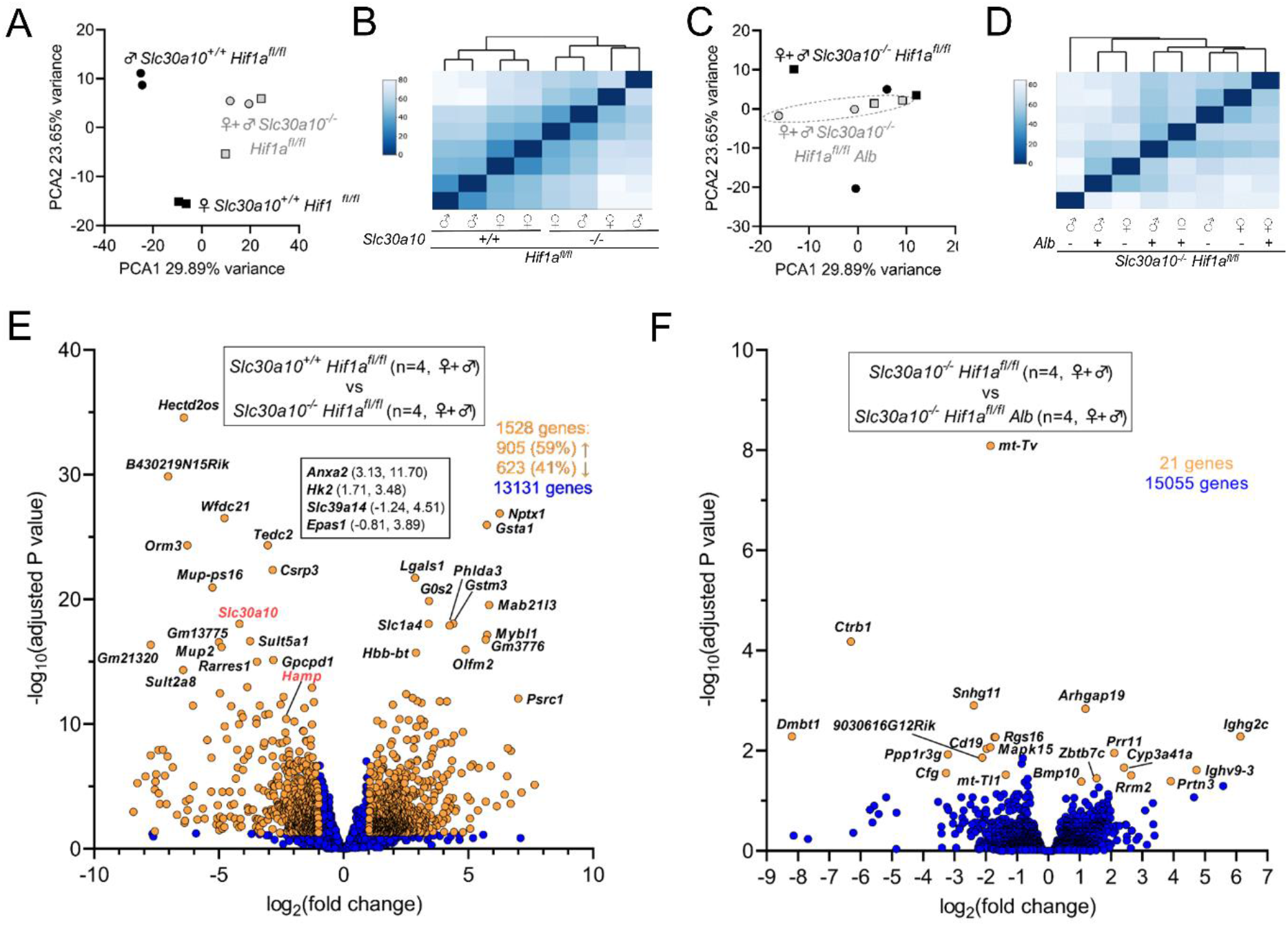
Hepatic Hif1a deficiency has minimal impact on hepatic gene expression in *Slc30a10*^*-/-*^ mice. Principal component (**A, B**) and similarity analysis (**C, D**) of samples and volcano plots (**E, F**) of genes differentially expressed between *Slc30a10*^*+/+*^ *Hif1a*^*fl/fl*^ and *Slc30a10*^*-/-*^ *Hif1a*^*fl/fl*^ mice (**A-C**) and between *Slc30a10*^*-/-*^ *Hif1a*^*fl/fl*^ and *Slc30a10*^*-/-*^ *Hif1a*^*fl/fl*^ *Alb* mice (**D-F**). Volcano plots represented as described in Fig. 6.

We next assessed the impact of Hif2a on differential gene expression in Slc30a10 deficiency by performing two comparisons: *Slc30a10*^*-/-*^ *Hif2a*^*fl/fl*^ versus *Slc30a10*^*+/+*^ *Hif2a*^*fl/fl*^ livers to identify genes differentially expressed in Slc30a10 deficiency in the *Slc30a10 Hif2a* cohort, and *Slc30a10*^*-/-*^ *Hif2a*^*fl/fl*^ *Alb* versus *Slc30a10*^*-/-*^ *Hif2a*^*fl/fl*^ livers to identify Hif2a-dependent genes differentially expressed in Slc30a10 deficiency. For the *Slc30a10*^*-/-*^ *Hif2a*^*fl/fl*^ versus *Slc30a10*^*+/+*^ *Hif2a*^*fl/fl*^ comparison, samples segregated by *Slc30a10* genotype and sex (Fig. 8A, B). For the *Slc30a10*^*-/-*^ *Hif2a*^*fl/fl*^ *Alb* versus *Slc30a10*^*-/-*^ *Hif2a*^*fl/fl*^ comparison, samples clustered by *Slc30a10* genotype and sex but less prominently than for the *Slc30a10*^*-/-*^ *Hif2a*^*fl/fl*^ versus *Slc30a10*^*+/+*^ *Hif2a*^*fl/fl*^ comparison (Fig. 8C, D). 1712 genes, including *Slc30a10* (downregulated in *Slc30a10*^*-/-*^ livers), *Serpine1, Hk2, Anxa2* (all upregulated), and *Hamp* (downregulated), were differentially expressed in *Slc30a10*^*-/-*^ *Hif2a*^*fl/fl*^ livers relative to *Slc30a10*^*+/+*^ *Hif2a*^*fl/fl*^ livers (Fig. 8E). 1041 genes, including *Anxa2, Serpine1* (both downregulated with Hif2a deficiency in *Slc30a10*^*-/-*^ livers), and *Hamp* (upregulated), were differentially expressed in *Slc30a10*^*-/-*^ *Hif2a*^*fl/fl*^ *Alb* vs. *Slc30a10*^*-/-*^ *Hif2a*^*fl/fl*^ livers (Fig. 8F). This analysis indicated that hepatic Hif2a deficiency had a prominent impact on differential gene expression in *Slc30a10*^*-/-*^ livers.

**Fig. 8:**
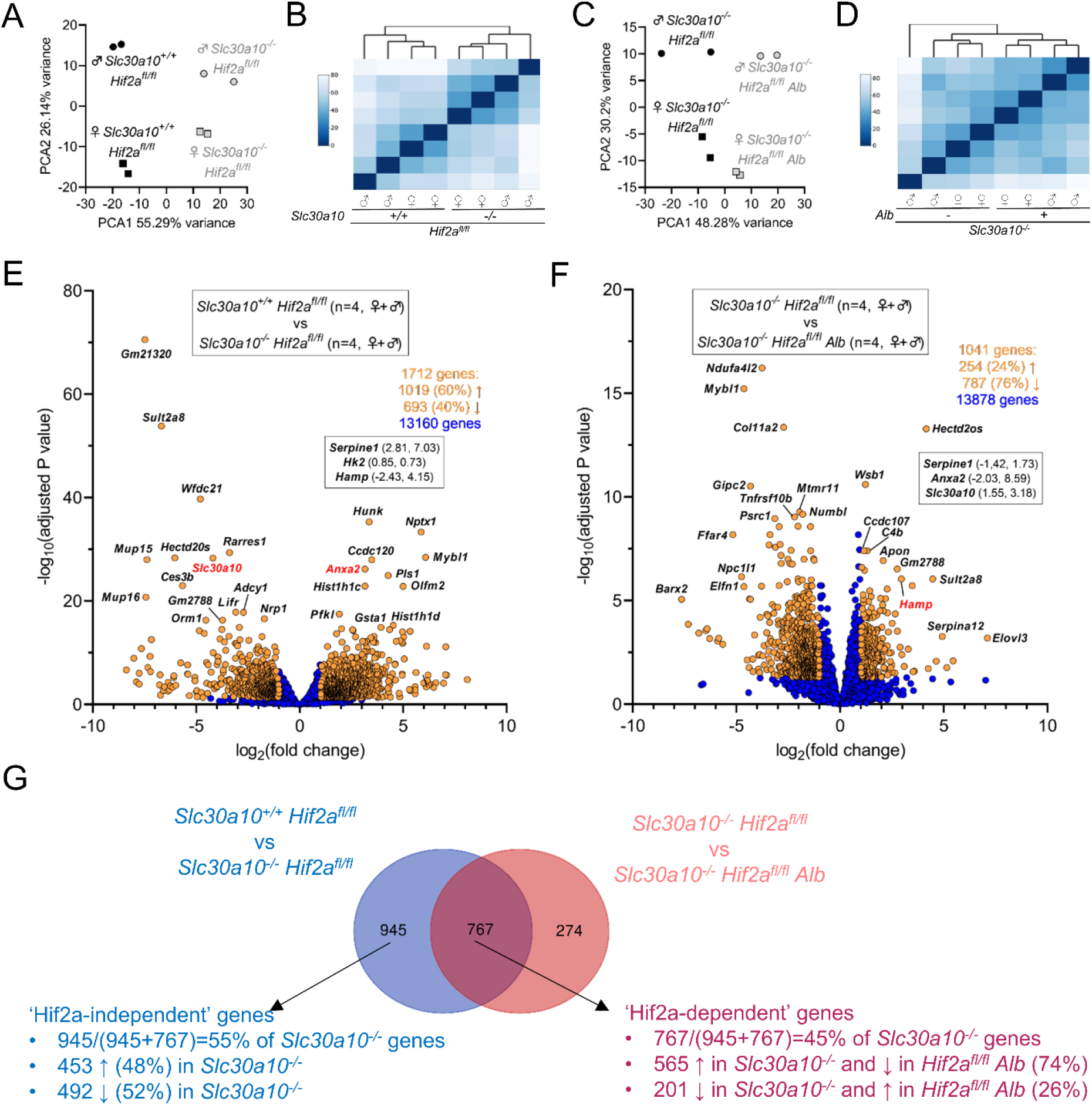
Hepatic Hif2a deficiency has a prominent impact on hepatic gene expression in *Slc30a10*^*-/-*^ mice. Principal component (**A, B**) and similarity analysis (**C, D**) of samples and volcano plots (**E, F**) of genes differentially expressed between *Slc30a10*^*+/+*^ *Hif2a*^*fl/fl*^ and *Slc30a10*^*-/-*^ *Hif2a*^*fl/fl*^ mice (**A-C**) and between *Slc30a10*^*-/-*^ *Hif2a*^*fl/fl*^ and *Slc30a10*^*-/-*^ *Hif2a*^*fl/fl*^ *Alb* mice (**D-F**). Volcano plots are represented as described in Fig. 6. (**G**) Venn diagram of differentially expressed genes from (**E, F**).

To further explore the impact of Hif2 on hepatic gene expression in Slc30a10 deficiency, we next assigned genes differentially expressed in *Slc30a10 Hif2a* mice to three groups (Fig. 8G). The first group, termed ‘Hif2a-independent’, consisted of 945 genes differentially expressed with Slc30a10 deficiency but not impacted by hepatic Hif2a deficiency. This group represented 55% of all genes differentially expressed with Slc30a10 deficiency. Within this group, 453 genes (48% of 945 genes) were upregulated with Slc30a10 deficiency and aligned largely with cell cycle (Fig. S7A-D). While these genes were denoted Hif2a-independent given their adjusted p values, expression of all of these genes decreased with Hif2a deficiency, albeit insignificantly (Fig. S7E). The remaining 492 genes (52% of 945 genes) in the Hif2a-independent group were downregulated with Slc30a10 deficiency and aligned mainly with metabolic processes (Fig. S8A-D). While these genes were denoted Hif2a-independent given their adjusted p values, expression of 95% of these genes increased with Hif2a deficiency, albeit insignificantly (Fig. S8E).

The second group of *Slc30a10 Hif2a* genes, termed ‘Hif2a-dependent’, consisted of 767 genes differentially expressed by Slc30a10 deficiency and then by hepatic Hif2a deficiency in Slc30a10-deficient mice (Fig. 8G). Within this group, 565 genes (74% of 767 genes) were upregulated with Slc30a10 deficiency then downregulated with Hif2a deficiency (Fig. S9). These genes aligned largely with cell cycle. The remaining 201 genes (26% of 7676 genes) were downregulated with Slc30a10 deficiency then upregulated with Hif2a deficiency (Fig. S10). These genes aligned mainly with metabolic processes.

The third group of *Slc30a10 Hif2a* genes were not differentially expressed by Slc30a10 deficiency but were differentially expressed by hepatic Hif2a deficiency in Slc30a10-deficient mice. This group consisted of 274 genes which aligned largely with response to stimuli (Fig. S11).

For the final analysis of our RNA-seq data, we compared gene expression in *Slc30a10*^*-/-*^ *Hif1a*^*fl/fl*^ *Alb* and *Slc30a10*^*-/-*^ *Hif2a*^*fl/fl*^ *Alb* livers. Samples clustered by sex and *Hif* genotype (Fig. 9A-B). 1468 genes were differentially expressed, including *Serpine1, Anxa2, Hk2, Epo* (all downregulated with Hif2a deficiency in *Slc30a10*^*-/-*^ livers) and *Hamp* (upregulated with Hif2a deficiency) (Fig. 9C). Most genes were aligned with cell cycle and metabolism (Fig. 9D, E). The majority of cell cycle-aligned genes were downregulated by hepatic Hif2a deficiency, while roughly equal numbers of metabolism-aligned were up- and downregulated by hepatic Hif2a deficiency (Fig. 9F).

**Fig. 9:**
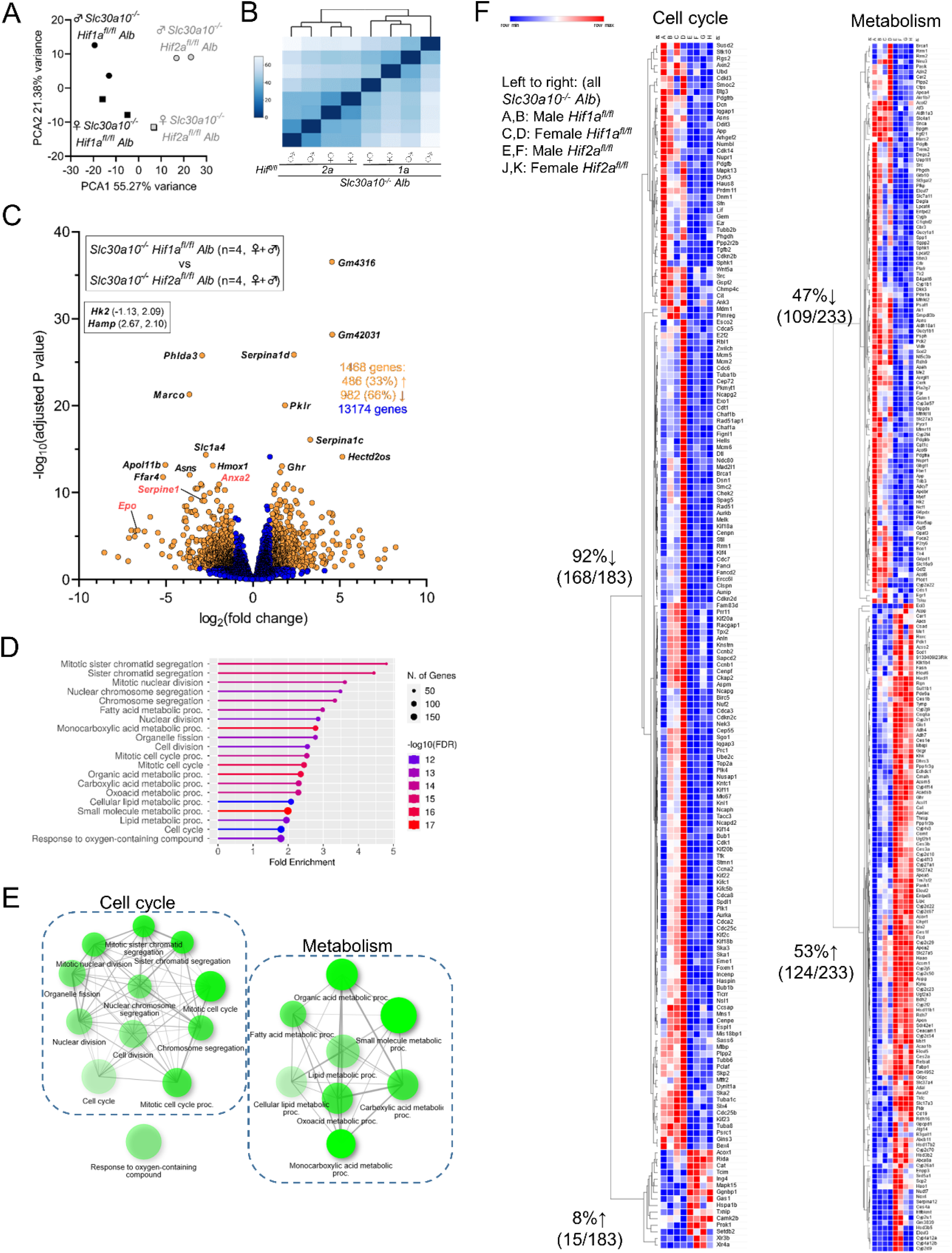
Hepatic genes differentially expressed between *Slc30a10*^*-/-*^ *Hif1a*^*fl/fl*^ *Alb* and *Slc30a10*^*-/-*^ *Hif2a*^*fl/fl*^ *Alb* mice align largely with cell cycle and metabolism pathways. (**A-C**) Principal component (**A**) and similarity analysis (**B**) of samples and volcano plot (**C**) of genes differentially expressed between *Slc30a10*^*-/-*^ *Hif1a*^*fl/fl*^ *Alb* and *Slc30a10*^*-/-*^ *Hif2a*^*fl/fl*^ *Alb* mice. Volcano plot is represented as in Fig. 6. (**D-E**) Gene enrichment analysis. (**F**) Heat map of gene expression for genes aligned with pathways encircled by dashed shape in (**E**).

We next considered the absence of *Epo* as a differentially expressed gene in several RNA-seq analyses, despite the prominent change in liver *Epo* RNA levels in *Slc30a10-/-* mice. For qPCR, samples with undetectable *Epo* levels were arbitrarily assigned Ct values of 40 for calculation of fold-differences in expression (Fig. S12). However, in RNA-seq analyses, *Epo* raw counts for most samples were below the cut-off of 10 and excluded from further analysis.

### Hepatic Hif2a deficiency impacts iron homeostasis in Slc30a10^-/-^ mice

RNA-seq analyses of *Slc30a10* livers also indicated that hepcidin (*Hamp*) was downregulated with Slc30a10 deficiency in a Hif2a-dependent manner. Hepcidin is a hormone synthesized mainly by the liver that inhibits dietary iron absorption and macrophage iron export (25). It is upregulated by iron excess and inflammation and downregulated by iron deficiency and in settings of increased iron demand such as Epo excess. We next investigated if the decreased hepcidin expression had functional impact on iron homeostasis in *Slc30a10*^*-/-*^ mice. We first validated decreased liver hepcidin RNA levels and serum hepcidin levels in *Slc30a10*^*-/-*^ mice (Fig. 10A, B). Consistent with hepcidin deficiency, *Slc30a10*^*-/-*^ mice exhibited increased absorption of gavaged ^59^Fe (Fig. 10C). Total and non-heme iron levels were decreased in *Slc30a10*^*-/-*^ spleen (Fig. 10D, E), also consistent with hepcidin deficiency—since hepcidin inhibits macrophage iron export, hepcidin deficiency leads to increased iron export from red pulp macrophages, leading to decreased spleen iron levels.

**Fig. 10:**
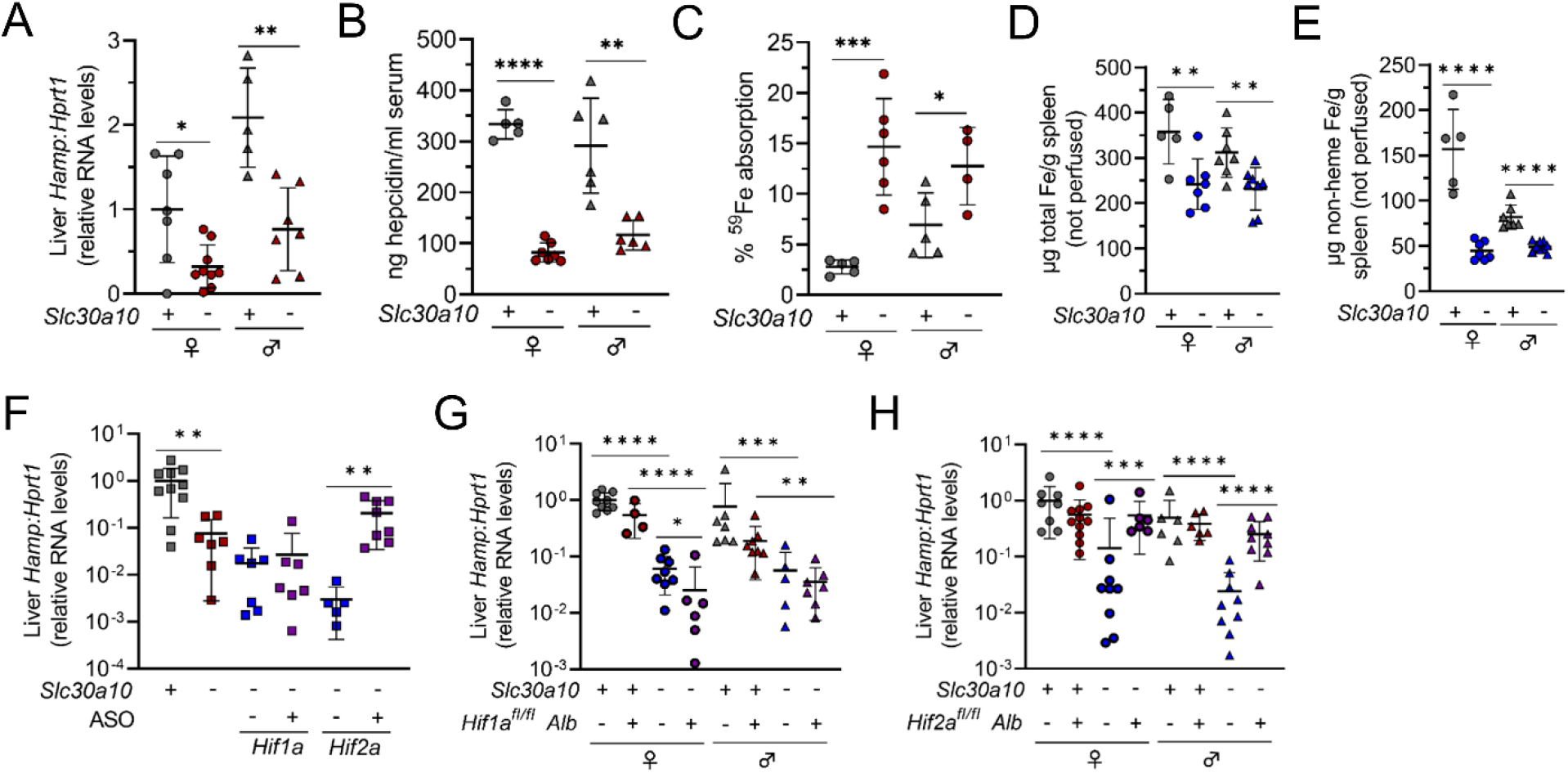
Hepatic Hif2a deficiency impacts iron (Fe) homeostasis in *Slc30a10*^*-/-*^ mice. (**A-D**) Mice from Fig. 1A-F were analyzed for: (**A**) liver hepcidin (*Hamp*) RNA levels by qPCR; (**B**) serum Epo levels by ELISA; (**C**) ^59^Fe absorption by gastric gavage; (**D**) total Fe levels in spleen by ICP-ES; (**E**) non-heme Fe levels in spleen by bathophenanthroline-based assay. (**F-H**) Mice from Fig. 2-4 were analyzed for liver *Hamp* RNA levels by qPCR. Data are represented and statistics performed and annotated as in Fig. 1.

Our RNA-seq analyses presented above indicated that *Hamp* is downregulated by Slc30a10 deficiency in a Hif2a-dependent manner. To validate and further explore the link between Hif2a and hepcidin, we measured liver hepcidin RNA by QPCR in *Slc30a10*^*-/-*^ mice with altered Hif levels. We first considered our ASO-treated *Slc30a10* cohort. We noted above that treatment of *Slc30a10*^*-/-*^ mice with *Hif2a* ASOs, but not *Hif1a* ASOs, decreased liver *Epo* RNA levels (Fig. 3F). Treatment with *Hif2a* ASOs, but not *Hif1a* ASOs, increased liver hepcidin RNA levels (Fig. 10F). We also noted above that hepatic Hif2a deficiency, but not Hif1a deficiency, corrected liver *Epo* excess (Fig. 4F, 5F). Hepatocyte Hif2a deficiency, but not hepatocyte Hif1a deficiency, increased liver hepcidin RNA levels in *Slc30a10*^*-/-*^ mice (Fig. 10G, H). Overall, this indicated that hepatic Hif2a is essential for hepcidin downregulation in *Slc30a10*^*-/-*^ mice.

## Discussion

SLC30A10 deficiency is the first known inherited cause of Mn excess. It is characterized by neurologic and liver dysfunction, polycythemia, and EPO excess. Neurologic and liver disease are attributed to Mn toxicity, while polycythemia is attributed to EPO excess. The cause of EPO excess in this disease has yet to be established. Based upon the data presented above, we propose the following model to explain the link between SLC30A10 deficiency and EPO excess (see graphical abstract). SLC30A10 deficiency impairs gastrointestinal Mn excretion, leading to systemic Mn excess. Excess Mn is imported into the liver (and pancreas and small and large intestines) by SLC39A14. (SLC39A14 deficiency is the second reported inherited cause of Mn excess. Unlike SLC30A10 deficiency, SLC39A14 deficiency does not produce liver dysfunction, polycythemia, or EPO excess.) Liver Mn excess leads to increased HIF2-dependent EPO expression. EPO excess results in polycythemia and suppresses liver hepcidin expression, leading to increased dietary iron absorption with the majority of excess iron consumed by erythropoiesis.

The observation that hepatic Hif2a deficiency attenuates *Epo* excess and polycythemia in *Slc30a10*^*-/-*^ mice is consistent with a previous study showing that hepatic *Epo* expression is regulated by Hif2, not Hif1a (26). However, our work raises several questions. First, why does hepatic Hif2a deficiency also attenuate Mn excess in *Slc30a10*^*-/-*^ mice? One possibility is that decreased tissue Mn levels reflect correction of polycythemia, but mice were perfused with saline prior to tissue harvest. Another possibility is that hepatic Hif2a deficiency increases Mn excretion. Bile Mn levels did not increase in *Slc30a10*^*-/-*^ mice with hepatic Hif2a deficiency, suggesting that increased biliary Mn excretion was not a factor. Urinary excretion is unlikely, as Mn excretion by the kidneys is traditionally viewed as minimal. Gastrointestinal Mn excretion independent of the hepatobiliary route is a possibility, but this would have to be Slc30a10-independent given that our studies were based in *Slc30a10*^*-/-*^ mice. Another possibility is decreased dietary Mn absorption. While HIF2a and SLC30A10 have no known roles in regulating Mn absorption, *Slc30a10*^*-/-*^ mice do exhibit hepcidin deficiency. Hepcidin inhibits dietary iron absorption by post-translationally down-regulating ferroportin, a transport protein essential for export of iron from enterocytes into blood. Several studies report that that ferroportin can also transport Mn (27–33), while some do not support this notion (34). If hepcidin deficiency leads to increased ferroportin-dependent Mn absorption, hepatic Hif2a deficiency could decrease Mn levels by correcting hepcidin deficiency and suppressing Mn absorption.

The second issue raised by our work is the minimal impact of hepatic Hif1a deficiency on *Slc30a10*^*-/-*^ phenotypes. This observation is striking given that Mn excess impacts HIF1 in cell lines, as described in the introduction. In cell culture, Mn stimulates HIF activity by inhibiting PHDs. If this same mechanism is active in *Slc30a10*^*-/-*^ mice, why did our RNA-seq analysis show minimal impact of hepatic Hif1a deficiency on gene expression in *Slc30a10*^*-/-*^ livers? Perhaps Hif1a levels are too low to be impacted by PHD inhibition, but this seems unlikely, as both *Hif1a* and *Hif2a* RNA levels were decreased in *Slc30a10*^*-/-*^ livers. (We did attempt immunoblots to measure Hif1a and Hif2a protein levels in mouse tissues, but results were inconsistent and difficult to reproduce.) Another possible contributing factor is that pathophysiology in *Slc30a10*^*-/-*^ mice is not mimicked by exposure of cell lines to Mn—even dietary Mn excess in wild-type mice cannot achieve the levels of tissue Mn excess observed in *Slc30a10*^*-/-*^ mice (6). Yet another possibility is that SLC30A10 plays Mn-independent roles and this contributes somehow to the HIF2-specific liver phenotype in SLC30A10 deficiency, but we have no data in support of this speculation at this time.

The third prominent topic at hand is the significance of aberrant expression of genes beyond *Epo* in livers of *Slc30a10*^*-/-*^ mice. The majority of upregulated genes aligned with cell cycle, while the majority of downregulated genes aligned with metabolic processes. Hepatic Hif2a deficiency attenuated almost half of differential gene expression, indicating that HIF2 is a key determinant of hepatic gene expression in SLC30A10 deficiency. Given that HIF2 stimulates gene expression, we propose that upregulation of cell cycle genes in *Slc30a10*^*-/-*^ mice is a direct result of increased HIF2 activity, while the downregulation of metabolic process genes is due to an indirect consequence of increased HIF2 activity. HIFs are well known to regulate cell proliferation (35, 36) and can also impact metabolism of carbohydrates and lipids, as observed in diseases such as cancer, diabetes, and non-alcoholic fatty liver disease (37–40). While the specific HIF-dependent pathways aberrantly expressed in *Slc30a10*^*-/-*^ livers were not surprising given the established roles of HIFs in health and disease, it is not clear at this time how aberrant HIF2-dependent gene expression outside of EPO contributes to the pathophysiology of SLC30A10 deficiency.

Based upon the data presented above, we propose that the prominent impact of hepatic Hif2a deficiency on Mn excess, *Epo* excess, polycythemia, and hepatic gene expression warrants an investigation into the long-term treatment of *Slc30a10*^*-/-*^ mice with agents that inhibit Hif2a expression and/or activity. One option is Hif2a *ASOs* as we employed above. Another option is the HIF2 inhibitor Belzutifan (41), already approved for cancer treatment in patients with mutations in VHL, an E3 ubiquitin ligase essential for HIF ubiquitination and degradation. Current treatment options for SLC30A10 deficiency include chelation and oral iron supplementation, with the latter proposed to act by outcompeting dietary Mn for absorption. Our studies suggest HIF2 antagonism as a novel therapeutic option for this disease. On a final note, a recent study reported that Mn excess stimulates HIF-dependent *SLC30A10* expression in a homeostatic response to Mn toxicity and that PHD inhibitors protect mice against Mn toxicity, presumably by upregulating SLC30A10-dependent Mn excretion (20). We propose that treatment of patients with Mn excess with HIF-modulating agents be considered carefully, as our results suggest that HIF2 inhibition, not stabilization, is warranted in SLC30A10 deficiency.

## Methods

### Care, generation, and treatment of mice

Mouse studies were approved by the Institutional Animal Care and Use Committee at Brown University. Mice were bred and maintained in the animal care facility at Brown University. Mice were group-housed in ventilated cage racks, maintained on a 12-hour light-dark cycle with controlled temperature and humidity, and provided standard chow (LabDiet 5010 containing 120 ppm Mn) and water *ad libitum* unless otherwise specified. Littermates of the same sex were randomly assigned to experimental groups. *Slc30a10*^*-/-*^ and *Slc30a10*^*lox/lox*^ mice were generated as previously described(6) and maintained by backcrossing onto C57BL/6N mice (Jackson Laboratory). *Slc30a10*^*+/+*^ and *Slc30a10*^*-/-*^ mice were generated by crossing *Slc30a10*^*+/-*^ mice. To raise mice on Mn-deficient diets, mice were weaned onto diets containing 1 or 100 ppm Mn (Envigo TD.140497, TD.140499). *Slc39a14*^*-/-*^ mice were generated as previously described(42) and maintained by backcrossing onto 129S6/SvEvTac mice (Taconic). To generate mice with Slc30a10 and Slc39a14 deficiency, *Slc30a10*^*+/-*^ and *Slc39a14*^*+/-*^ mice were crossed, then *Slc30a10*^*+/-*^ *Slc39a14*^*+/-*^ progeny were bred together. To generate *Slc30a10*^*-/-*^ mice with hepatocyte Hif1a or Hif2a deficiency, *Slc30a10*^*+/-*^ were bred to mice expressing an albumin promotor-driven Cre recombinase transgene (B6N.Cg.*Speer6-ps1*^*Tg(Alb-Cre)21Mgn*^/J; “*Alb”*; Jackson Laboratory #018961) and B6.129-*Hif1a*^*tm3Rsjo*^/J (“*Hif1a*^*fl/fl*^”; Jackson Laboratory, #007561) or *Epas1*^*tm1Mcs*^*/J* (“*Hif2a*^*fl/fl*^”; Jackson Laboratory, #008407) mice in a multi-step breeding scheme. Mice for analysis were ultimately generated by crossing *Slc30a10*^*+/-*^ *Hif1a*^*fl/fl*^ and *Slc30a10*^*+/-*^ *Hif1a*^*+/fl*^ *Alb* mice and *Slc30a10*^*+/-*^ *Hif2a*^*fl/fl*^ and *Slc30a10*^*+/-*^ *Hif2a*^*+/fl*^ *Alb* mice. For ASO-based studies, *Slc30a10*^*+/+*^ mice were treated with sterile-filtered PBS and *Slc30a10*^*-/-*^ mice were treated with sterile-filtered GalNAc-control, *Hif1a*, or *Hif2a* ASO (10 µl/gram body weight, stock 0.25 mg/ml, Ionis Pharmaceuticals). ASOs were administered by intraperitoneal injection twice a week beginning at weaning until six weeks of age for a total of 12 doses. Animals were harvested two days after the last dose.

### Bile, blood, and tissue collection

For mice from which bile was not collected (Fig. 1, S1, 5a-d), mice were anesthetized by intraperitoneal injection of ketamine and xylazine. Blood was then collected by retro-orbital puncture into EDTA-coated tubes (BD) using heparinized capillary tubes (Fisher) then into serum collection tubes (BD) using nonheparinized capillary tubes (Fisher). Mice were euthanized by cervical dislocation and tissues collected for metal, DNA, RNA, and protein analysis. (Gastrointestinal tracts were washed of lumenal contents.)

For all other mice, mice were anesthetized with isoflurane then underwent bile, blood, and tissue collection. Bile was collected surgically by ligation of the common bile duct, cannulation of the gallbladder, and collection over 60 minutes as previously described(6). Bile volumes were measured every five minutes to permit calculation of bile flow rates. Blood was then collected by cardiac puncture, and mice were transcardially perfused with PBS to remove blood from tissues. Tissues were then collected and processed as above.

### Blood analysis

Complete blood counts were performed on freshly isolated anticoagulated blood using VetAbcPlus (Sci). Serum Epo and hepcidin levels were measured using Mouse Erythropoietin/EPO Quantikine ELISA Kit (R&D) and Hepcidin Murine Compete ELISA Kit (Intrinsic Life Sciences) respectively.

### RNA analysis

For PCR array analysis, total RNA was extracted using Aurum Total RNA Mini Kit (BioRad). Concentration and purity of extracted RNA was determined by NanoDrop ND-1000 (Thermo Scientific, USA). Ribosomal RNA integrity was assessed by Agilent Bioanalyser. RNA (1 µg) was reverse transcribed to cDNA using iScript Advanced cDNA synthesis kit (BioRad). Each 96-well hypoxia array (BioRad) contained primers for 91 hypoxia signaling pathway-related genes and five endogenous controls for amplification, DNA contamination, reverse transcription, and RNA quality. qPCR was carried out on Viia 7 Real-Time PCR System (Applied Biosystems) using SsoAdvanced Universal SYBR Green Supermix (BioRad). Data were exported to BioRad CFX Manager Software and analyzed using relative quantification (2^ΔΔCt^) approach. A two-fold change in gene expression was used as a cut-off for up- or down-regulated genes.

For gene-specific qPCR, 100 to 200 mg tissue was homogenized in TRIzol (ThermoFisher) using 0.5 mm zirconium beads and Bullet Blender (Next Advance), followed by chloroform extraction, isopropanol precipitation, and 70% ethanol wash, as previously described(6). Standards were made by serially diluting mixtures of control and experimental samples, then processed identically as experimental samples. Samples underwent DNase treatment and cDNA synthesis using the High Capacity cDNA Reverse Transcription Kit with RNase Inhibitor (ThermoFisher). qPCR was performed using PowerUP SYBR Green Master Mix (ThermoFisher). Primers listed below were designed using Primer3 and Geneious. Amplicon fidelity was confirmed by Sanger sequencing. Primer concentrations were optimized by testing a series of combination of forward and reverse primers followed by melt curve analyses.

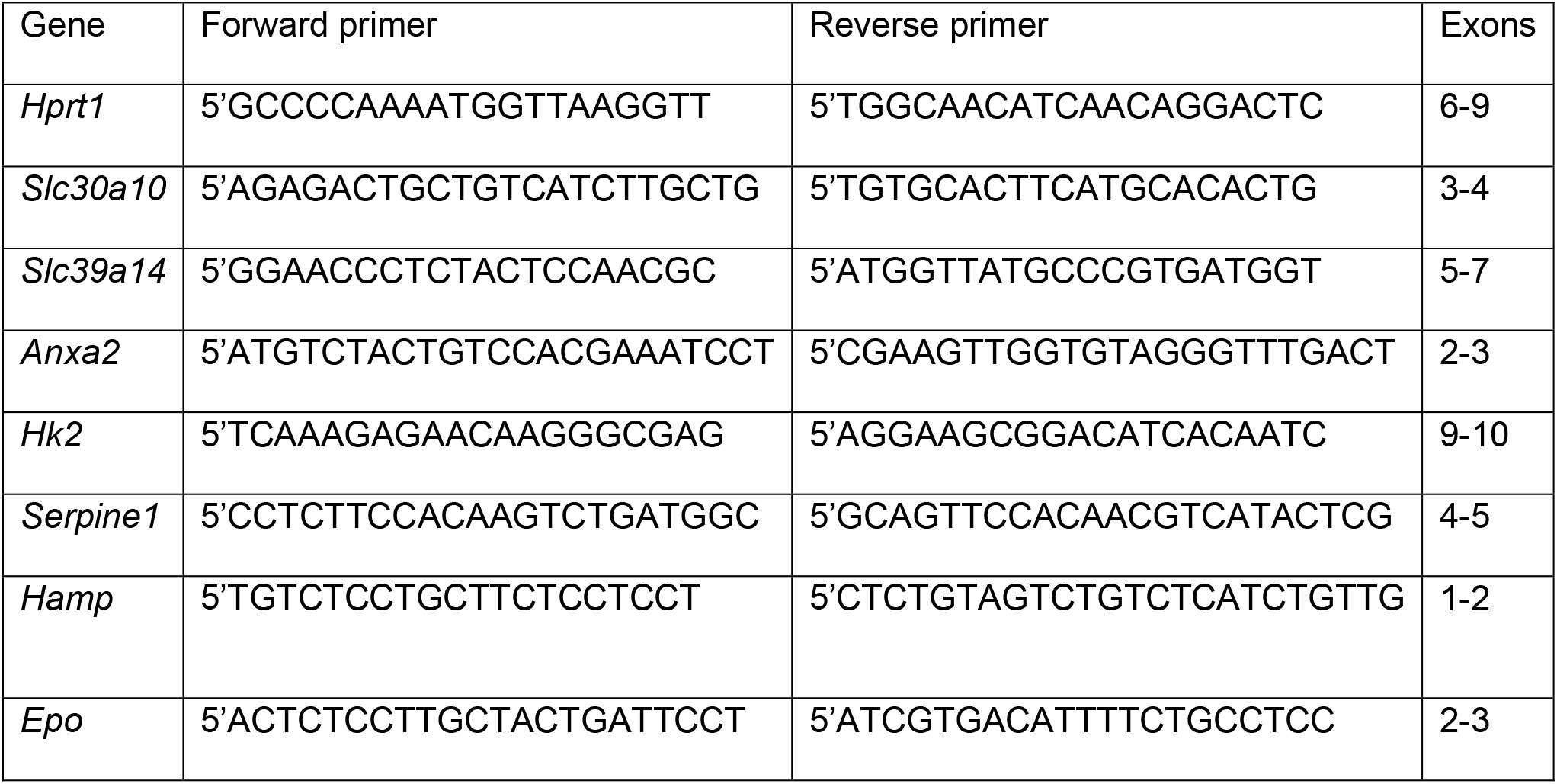
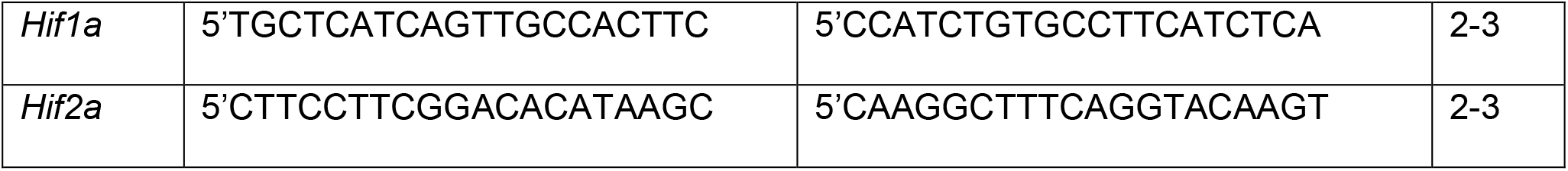

RNA extraction, library preparation, sequencing and analysis was conducted at Azenta Life Sciences (South Plainfield, NJ, USA) as follows. Total RNA was extracted from fresh frozen tissue samples using Qiagen RNeasy Plus Universal mini kit following manufacturer’s instructions (Qiagen, Hilden, Germany). RNA samples were quantified using Qubit 2.0 Fluorometer (Life Technologies, Carlsbad, CA, USA) and RNA integrity was checked using Agilent TapeStation 4200 (Agilent Technologies, Palo Alto, CA, USA). RNA sequencing libraries were prepared using the NEBNext Ultra II RNA Library Prep for Illumina using manufacturer’s instructions (NEB, Ipswich, MA, USA). Briefly, mRNAs were initially enriched with Oligod(T) beads. Enriched mRNAs were fragmented for 15 minutes at 94 °C. First strand and second strand cDNA were subsequently synthesized. cDNA fragments were end repaired and adenylated at 3’ends, and universal adapters were ligated to cDNA fragments, followed by index addition and library enrichment by PCR with limited cycles. The sequencing libraries were validated on the Agilent TapeStation (Agilent Technologies, Palo Alto, CA, USA), and quantified by using Qubit 2.0 Fluorometer (Invitrogen, Carlsbad, CA) as well as by quantitative PCR (KAPA Biosystems, Wilmington, MA, USA). The sequencing libraries were clustered on a flowcell. After clustering, the flowcell was loaded on the Illumina instrument (4000 or equivalent) according to manufacturer’s instructions. The samples were sequenced using a 2×150bp Paired End (PE) configuration. Image analysis and base calling were conducted by the Control software. Raw sequence data (.bcl files) generated the sequencer were converted into fastq files and de-multiplexed using Illumina’s bcl2fastq 2.17 software. One mismatch was allowed for index sequence identification. After investigating the quality of the raw data, sequence reads were trimmed to remove possible adapter sequences and nucleotides with poor quality. The trimmed reads were mapped to the reference genome available on ENSEMBL using the STAR aligner v.2.5.2b. The STAR aligner is a splice aligner that detects splice junctions and incorporates them to help align the entire read sequences. BAM files were generated as a result of this step. (BAM files were uploaded on the NCBI Sequence Read Archive under accession PRJNA924303.) Unique gene hit counts were calculated by using feature Counts from the Subread package v.1.5.2. Only unique reads that fell within exon regions were counted. After extraction of gene hit counts, the gene hit counts table was used for downstream differential expression analysis. Using DESeq2, a comparison of gene expression between the groups of samples was performed. The Wald test was used to generate p-values and Log2 fold changes. Genes with adjusted p-values < 0.05 and absolute log2 fold changes > 1 were called as differentially expressed genes for each comparison. A PCA analysis was performed using the “plotPCA” function within the DESeq2 R package. The top 500 genes, selected by highest row variance, were used to generate PCA plots. Venn diagrams were generated using bioinformatics.psb.ugent.be/webtools/Venn/. Gene ontology was performed using ShinyGO at http://bioinformatics.sdstate.edu/go/(43). Heatmaps on rlog-transformed gene counts were generated using Morpheus at https://software.broadinstitute.org/morpheus/.

### Metal measurements

For measurement of tissue total metal levels, 10-200 mg tissue were digested in 1000 µl 70% trace-metal-grade nitric acid (Fisher) at 65°C for two hours then diluted 25-fold in MilliQ water (Millipore Sigma) and analyzed by ICPES (ThermoScientific iCAP 7400 DUO) or GAAS (Perkin Elmer AAnalyst 600). (If tissues were isolated from perfused mice, tissues were lyophilized for 48 hours using LABCONCO FreeZone 6 Liter Freeze Dry System prior to acid digestion.) For measurement of blood metal levels, samples were digested with two volumes nitric acid at 65°C for two hours, diluted 25-fold with water, then analyzed by GFAAS. For measurement of bile metal levels, samples were digested twice with one volume nitric acid at 65°C until dry, twice with one volume hydrogen peroxide at 65°C until dry, resuspended in 2% nitric acid, then analyzed by GFAAS. For ICPES, a series of standards were analyzed and sample values were extrapolated from the generated curve. A quality control standard (IV-28, Inorganic Ventures) was run every ten samples to assess changes in sensitivity. If tissue size was small or metal levels too low, GFAAS was used. For GFAAS, standards were measured to create a standard curve. To correct for variations in sensitivity, a quality control standard was analyzed every ten samples (NIST, SRM 1640a). Irrespective of instrumentation, a correction curve was calculated based on quality control analysis and correction factor applied to each sample to control for changes in instrument sensitivity.

For measurement of tissue non-heme iron levels, 10-200 mg tissue was digested in 1 ml 3 N hydrochloric acid (Fisher)/10% trichloroacetic acid (Millipore Sigma) at 65 °C for two days, with 30 minutes vortexing each day, followed by centrifugation. Iron levels were measured by mixing 10 μl supernatants with 200 μl chromagen (five volumes MilliQ water; five volumes saturated sodium acetate (Fisher); one volume chromagen stock, consisting of 0.1% bathophenanthroline sulfonate (Millipore Sigma) and 1% thioglycolic acid (Millipore Sigma)) in a 96-well plate. Iron standards (Fisher) were included. After a ten-minute incubation, absorbances were measured at 535 nm. Mock digests without samples were included for this and all other metal analyses.

### ^59^Fe absorption studies

Mice were fasted in metabolic cages (Tecniplast) for four hours to clear the upper gastrointestinal (GI) tract of most contents prior to gavage. To assess iron absorption, each mouse was gavaged with 1 µCi ^59^FeCl_2_ (Perkin Elmer) in 2.21 mM FeCl_3_ in 1 M ascorbic acid filter-sterilized. After gavage, mice were returned to the metabolic cage and harvested after one hour. All mice were then euthanized by cervical dislocation and GI tract from stomach to rectum as well as gallbladder were removed from each mouse. Radioactivity levels were measured using a Triathler Gamma Counter with external NaI well-type crystal detector (Hidex). All samples were counted for 15 seconds. Whole GI tract with contents was counted first in a 15 mL conical tube with stomach positioned closest to the detector. Gallbladder was counted in a 1.5 mL centrifuge tube. Body levels were measured with mice positioned headfirst in a 50 mL conical. After measuring levels in whole GI tract with contents, the whole GI tract was separated into stomach, small intestine, cecum, and large intestine. Each compartment was cleaned out and rinsed. Radioactivity for each rinsed compartment was measured by placing tissue in 1.5 mL centrifuge tube. Percent absorption was calculated as sum of radioactivity in whole carcass/total radioactivity.

### Statistics (for all non-RNA-seq analyses)

Statistics were performed using GraphPad Prism 9. Data were tested for normal distribution by Shapiro-Wilk test. If not normally distributed, data were log transformed. Groups within each sex were compared by one-way ANOVA with Tukey’s multiple comparisons test or by unpaired, two-tailed t test as indicated in figure legends. P<0.05 was considered significant. Data are represented as means +/- standard deviation.

## Author contributions

Study design: MP, SG, MA, TBB; execution of experiments and data analysis: MP, JZ, CJM, HLK, BD, JAA, TBB; provision of reagents: SG, MA; manuscript writing: TBB; manuscript revision: all authors.

## Acknowledgements

Funding: NIH DK84122 (T.B.B.), DK110049 (T.B.B.), DK117524 (C.J.M.), ES007272-24 T32 (C.J.M.), GM077995 T32 (H.L.C.). We thank Christoph Schorl and the Genomics Core for assistance with qPCR, and Joseph Orchardo, David Murray, and Marcelo da Rosa Alexandre for assistance with metal analysis.

