## Supplemental figures for "Hypoxia-inducible factor 2 is a key determinant of manganese excess and polycythemia in SLC30A10 deficiency"

[&]Currently at Myovant Sciences, Brisbane, CA, 94005

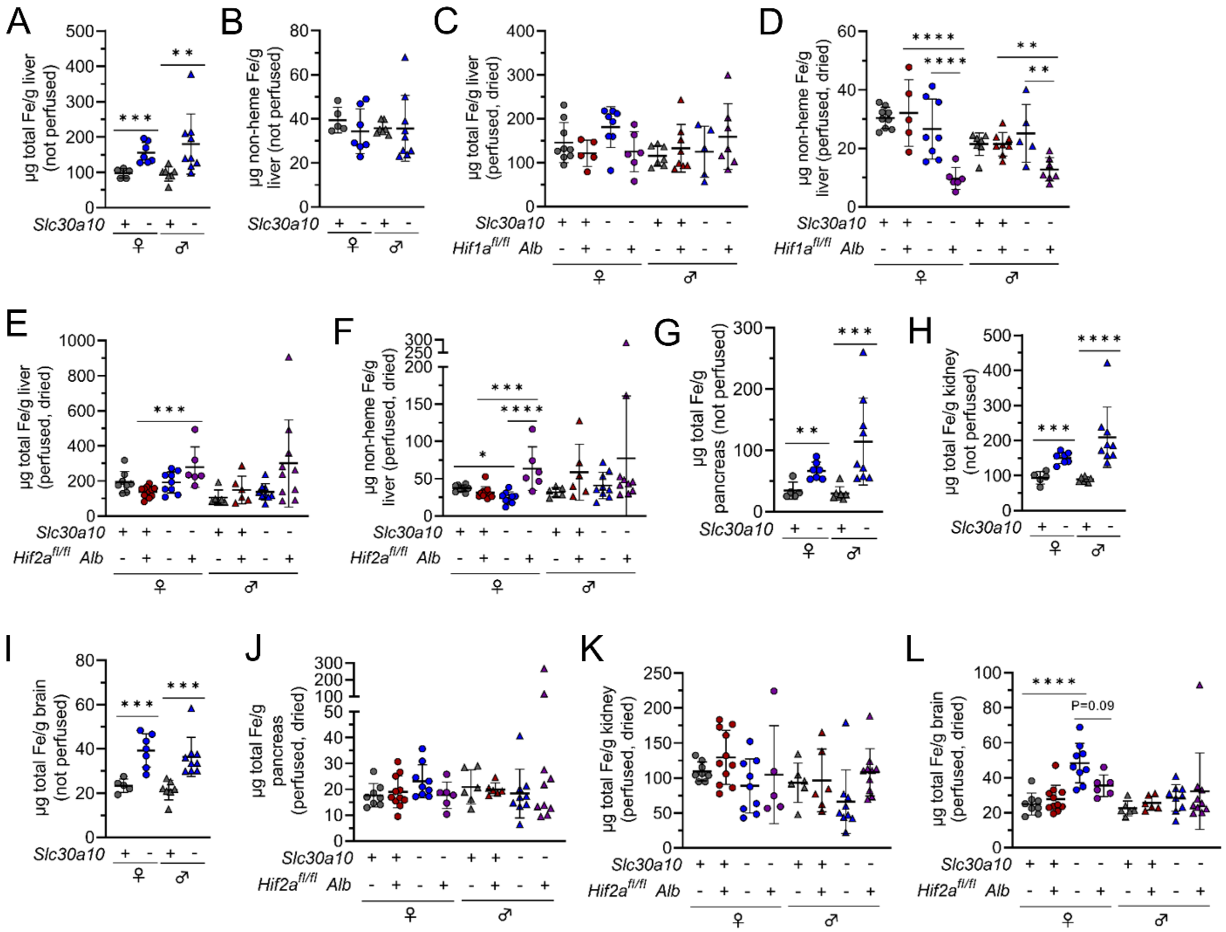


**Fig. S1: Perfusion of mice prior to tissue harvest impacts tissue iron (Fe) levels.** (**A**) Total Fe levels in livers from two-month-old *Slc30a10*$^{+/+}$ and *Slc30a10*$^{-/-}$ mice, measured by ICP-ES and previously published (6), are reproduced here for reference. (**B**) Non-heme Fe levels were measured by bathophenanthroline-based assay in livers from two-month-old *Slc30a10*$^{+/+}$ and *Slc30a10*$^{-/-}$ mice described in (6). (**C-F**) Total (**C, E**) and non-heme (**D, F**) Fe levels were measured in two-month-old *Slc30a10 Hif1a* (**C, D**) and *Slc30a10 Hif2a* (**E, F**) mice first described in Fig. 4 and 5. (**G-I**) Total Fe levels in pancreas (**G**), kidney (**H**), and brain (**I**) from two-month-old *Slc30a10*$^{+/+}$ and *Slc30a10*$^{-/-}$ mice, measured by ICP-ES and previously published (6), are reproduced here for reference. (**J-L**) Total Fe levels in pancreas (**J**), kidney (**K**), and brain (**L**) from *Slc30a10 Hif2a* mice measured by ICP-ES. Note that mice represented in (**A, B, G-I**) were not perfused with saline prior to tissue harvest; for all other panels, mice were perfused with saline and tissues dried prior to metal analysis. Data are represented and statistics performed and annotated as in Fig. 1.

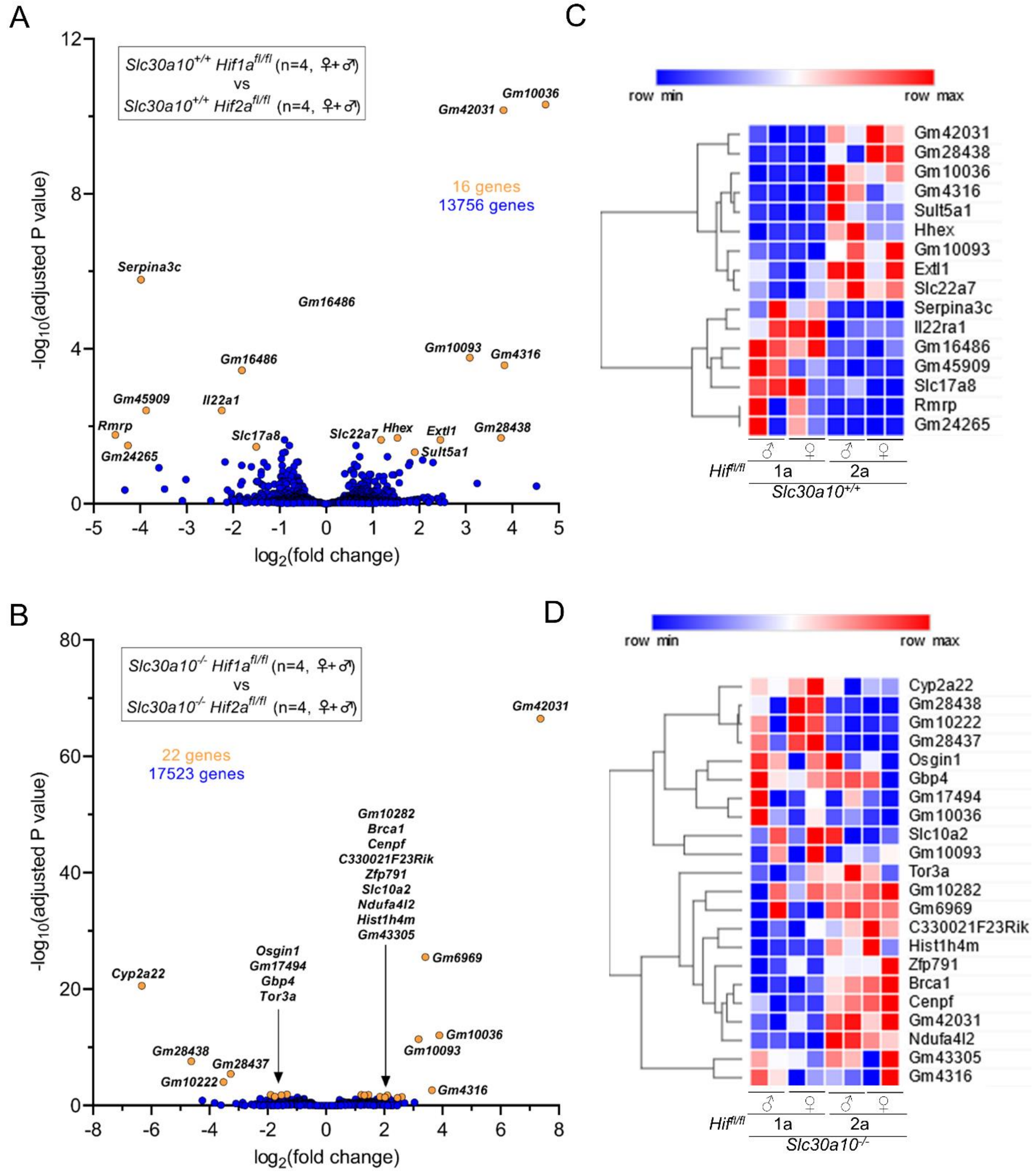


**Fig. S2: Floxed *Hif1a* and *Hif2a* alleles do not have prominent impact on hepatic gene expression.** Volcano plots (**A, B**) and heat maps (**C, D**) of genes differentially expressed between *Slc30a10*$^{+/+}$ *Hif1a*$^{fl/fl}$ and *Slc30a10*$^{+/+}$ *Hif2a*$^{fl/fl}$ livers (**A, C**) and between *Slc30a10*$^{-/-}$ *Hif1a*$^{fl/fl}$ and *Slc30a10*$^{-/-}$ *Hif2a*$^{fl/fl}$ livers (**B, D**). Volcano plots are represented as in Fig. 6.

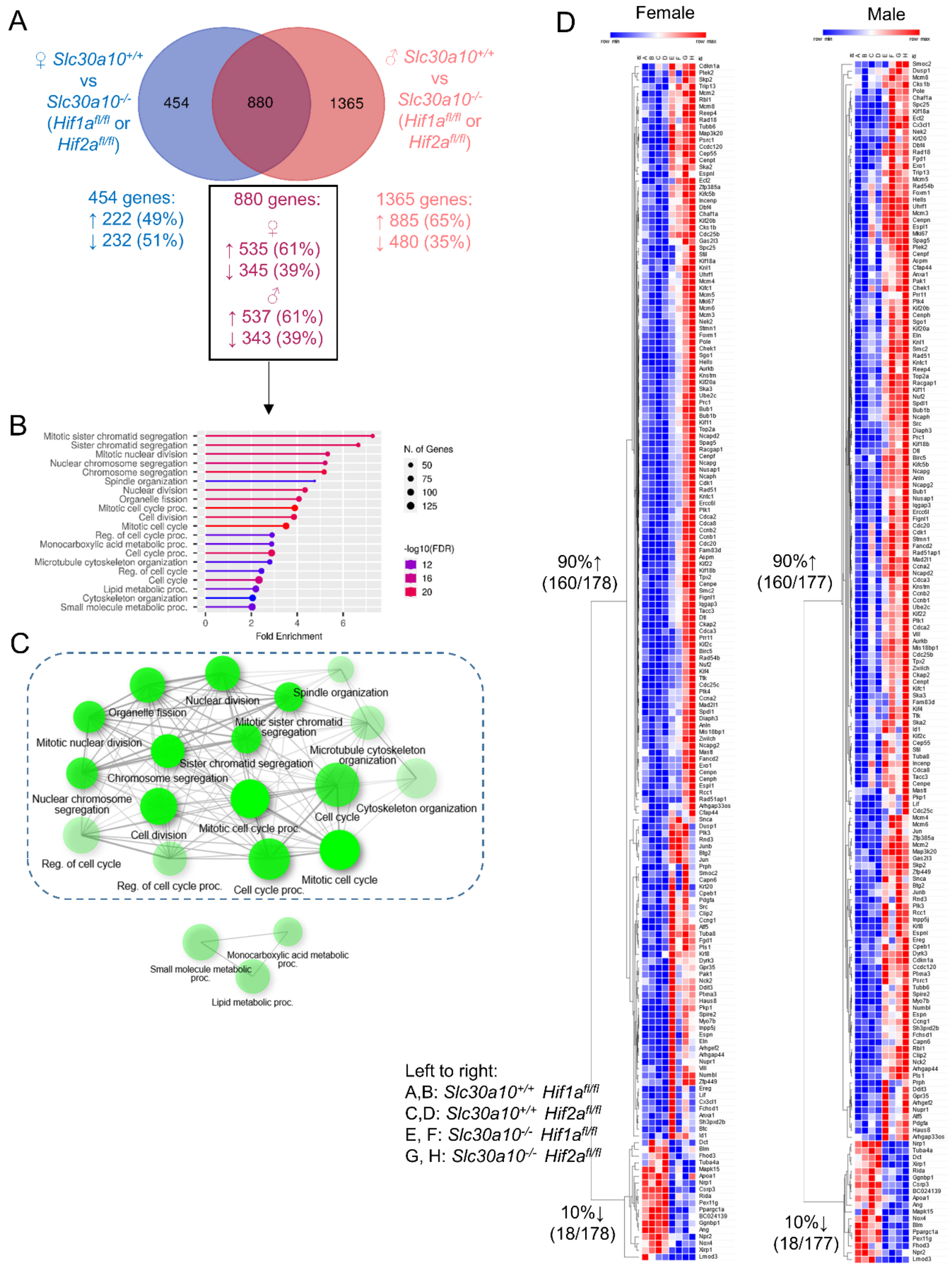


**Fig. S3: Hepatic genes differentially expressed in both female and male *Slc30a10$^{-/-}$* mice align largely with metabolic processes.** (**A**) Venn diagram of differentially expressed genes from Fig. 6. (**B, C**) Gene enrichment analysis. (**D**) Heat map of differentially expressed genes aligned with pathways encircled by dashed line in (**C**).

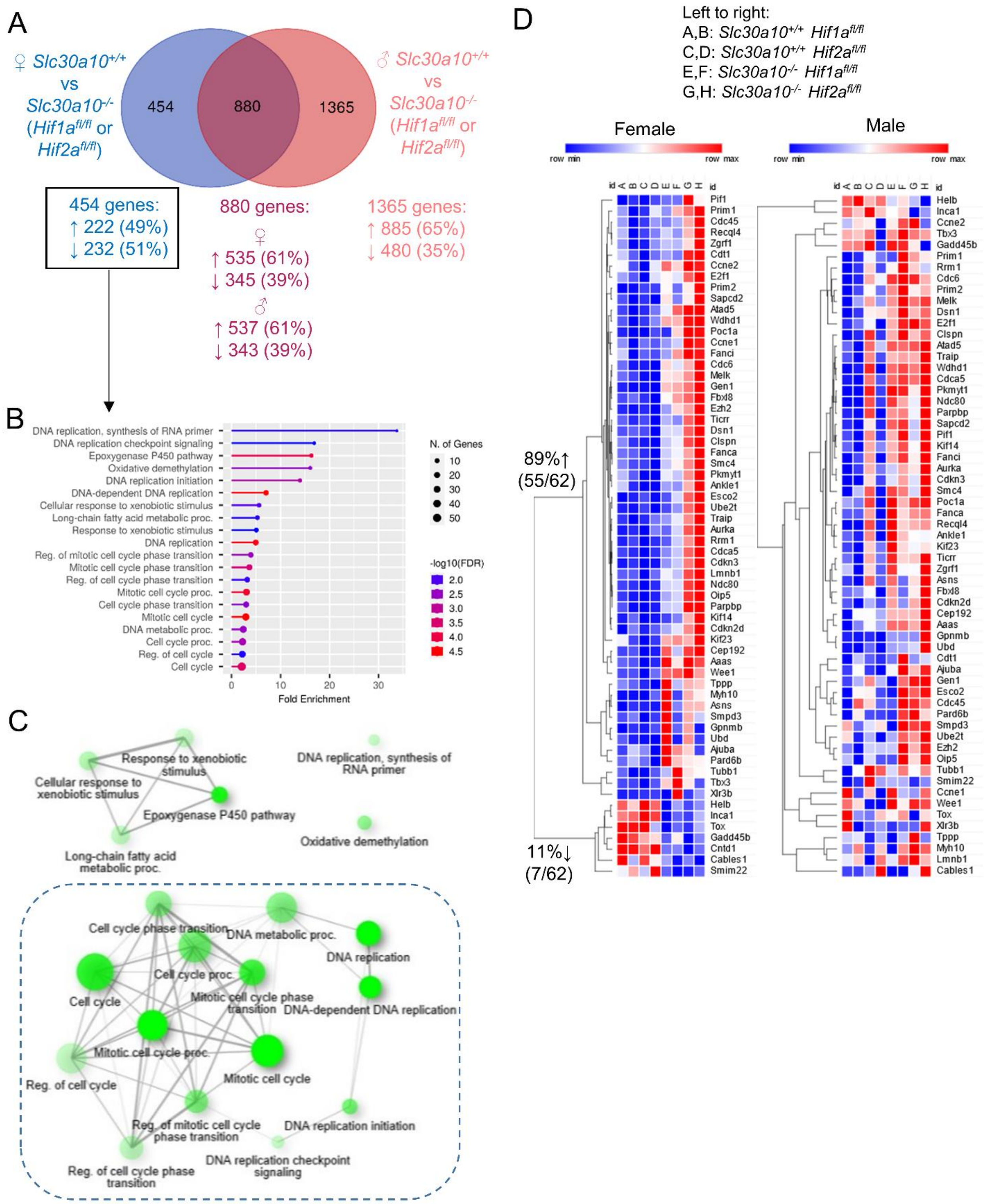


**Fig. S4: Hepatic genes differentially expressed in female but not male *Slc30a10*$^{-/-}$ mice align largely with cell cycle and DNA replication.** (**A**) Venn diagram of differentially expressed genes from Fig. 6. (**B, C**) Gene enrichment analysis. (**D**) Heat map of differentially expressed genes aligned with pathways encircled by dashed line in (**C**).

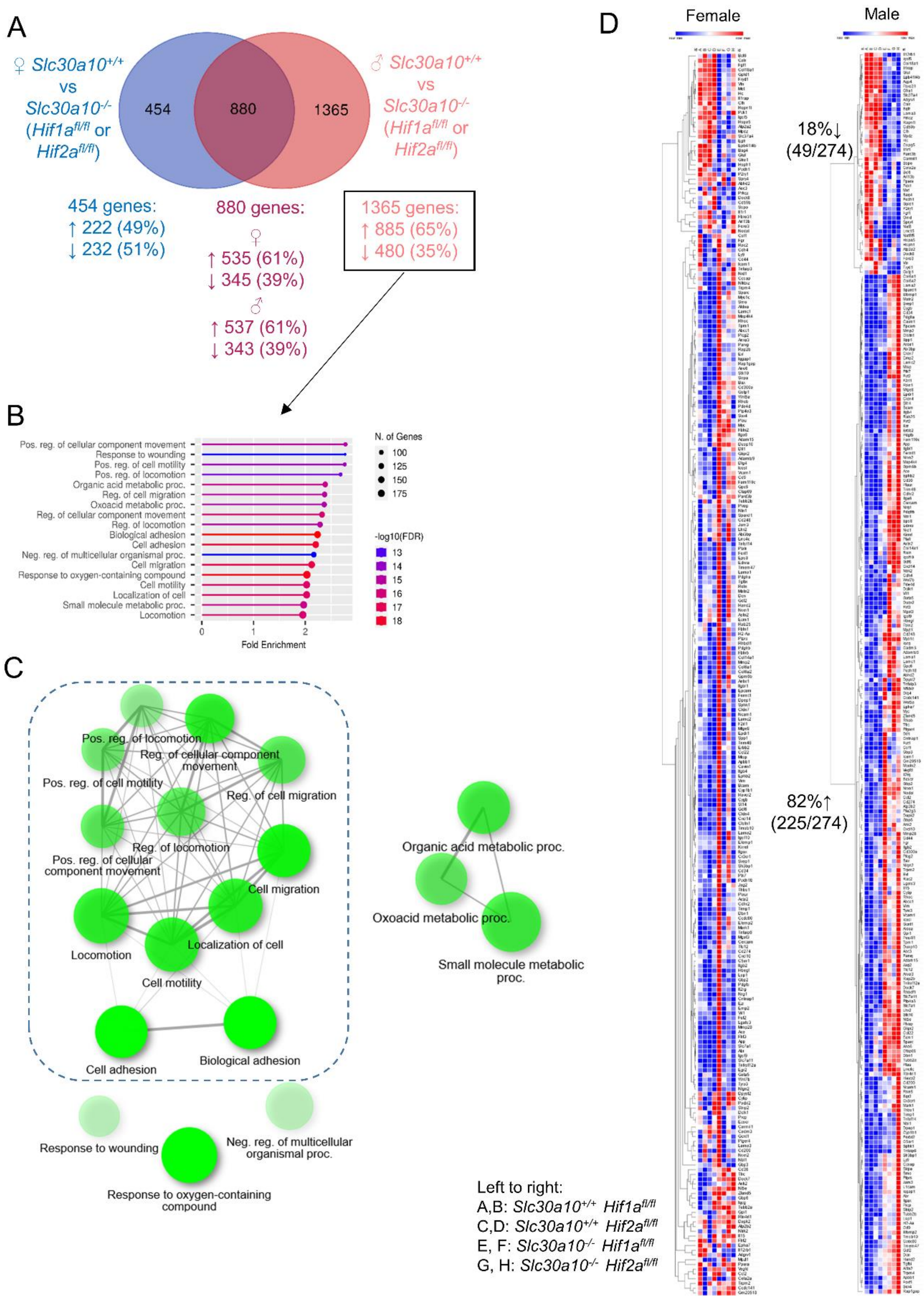


**Fig. S5: Hepatic genes differentially expressed in male but not female *Slc30a10*$^{-/-}$ mice align largely with cell motility.** (**A**) Venn diagram of differentially expressed genes from Fig. 6. (**B, C**) Gene enrichment analysis. (**D**) Heat map of differentially expressed genes aligned with pathways encircled by dashed line in (**C**).

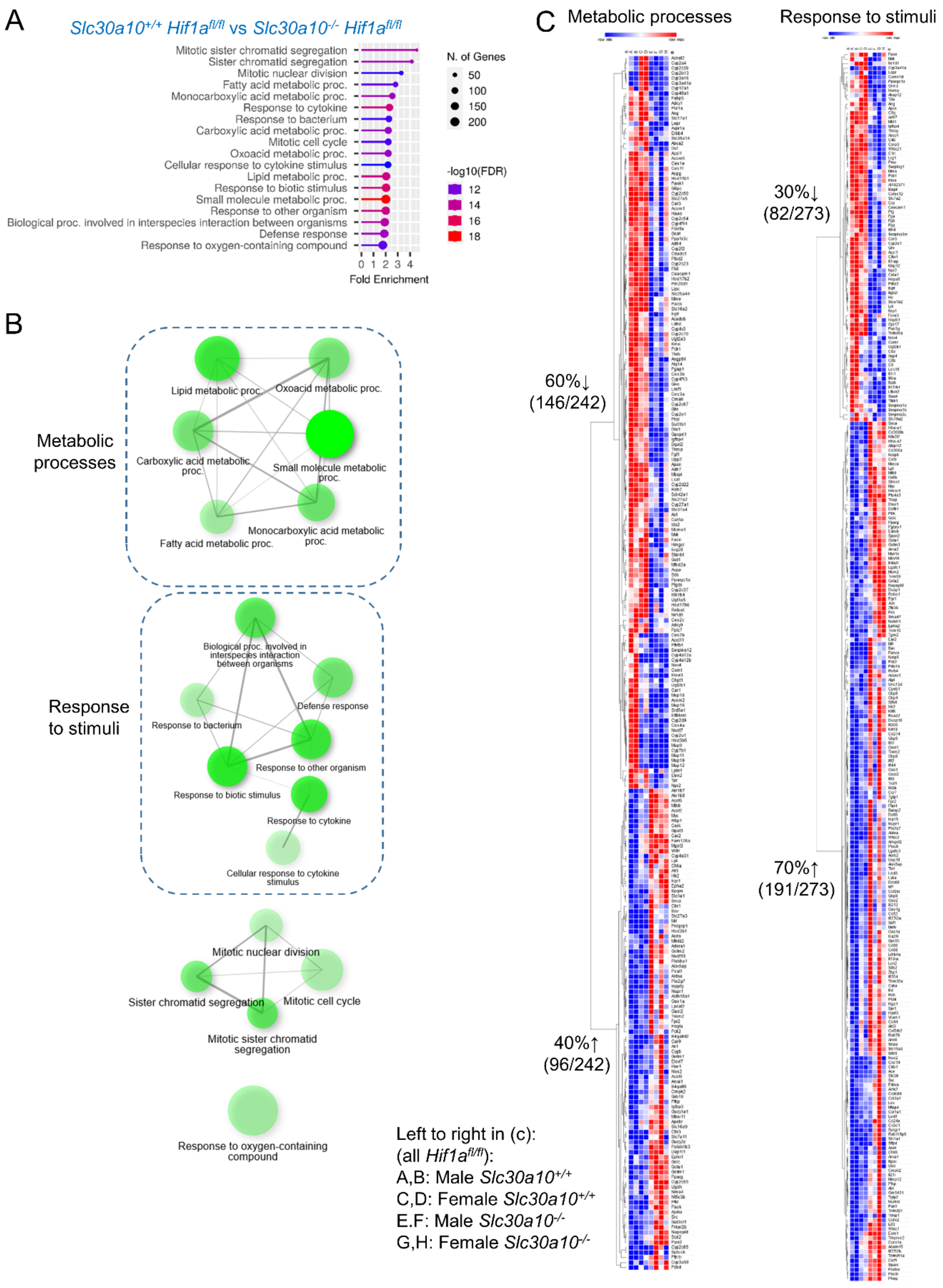


**Fig. S6: Hepatic genes differentially expressed in *Slc30a10*$^{-/-}$ *Hif1a*$^{fl/fl}$ mice, relative to *Slc30a10*$^{+/+}$ *Hif1a*$^{fl/fl}$ mice, align largely with metabolic processes and response to stimuli.** (**A, B**) Gene enrichment analysis. (**C**) Heatmap of differentially expressed genes aligned with pathways encircled by dashed lines in (**B**).

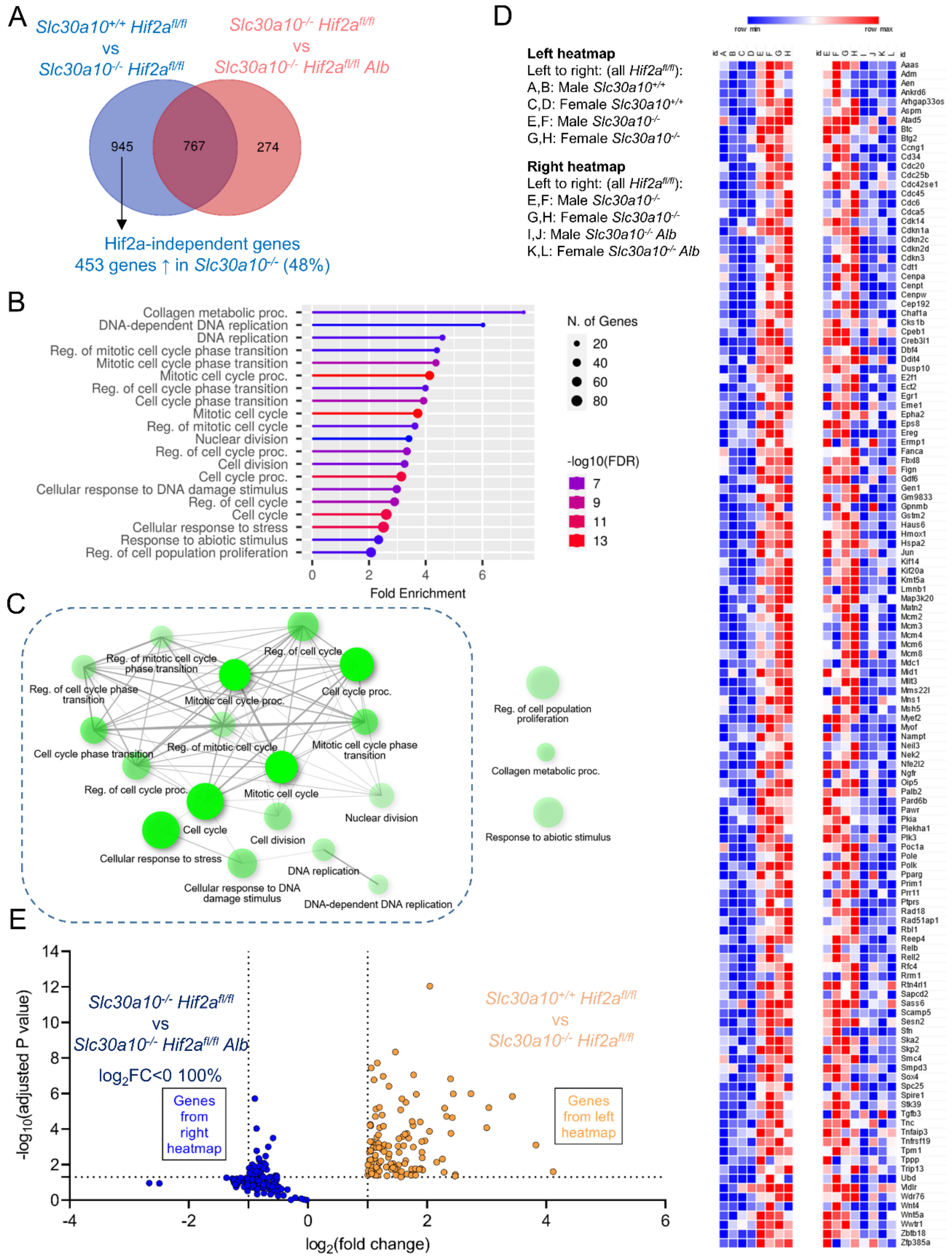


**Fig. S7: Hepatic genes upregulated in *Slc30a10*$^{-/-}$ mice but not impacted by hepatic Hif2a deficiency align largely with cell cycle.** (**A**) Venn diagram partially reproduced from Fig. 8G. (**B, C**) Gene enrichment analysis. (**D**) Heat map of differentially expressed genes aligned with pathways encircled by dashed line in (**C**). (**E**) Volcano plot of genes from left heatmap from (**D**) plotted as orange circles and from right heatmap from (**D**) as blue circles, indicating that all genes not significantly impacted by hepatic Hif2a deficiency are downregulated, i.e. $\log_2$(fold change)<0.

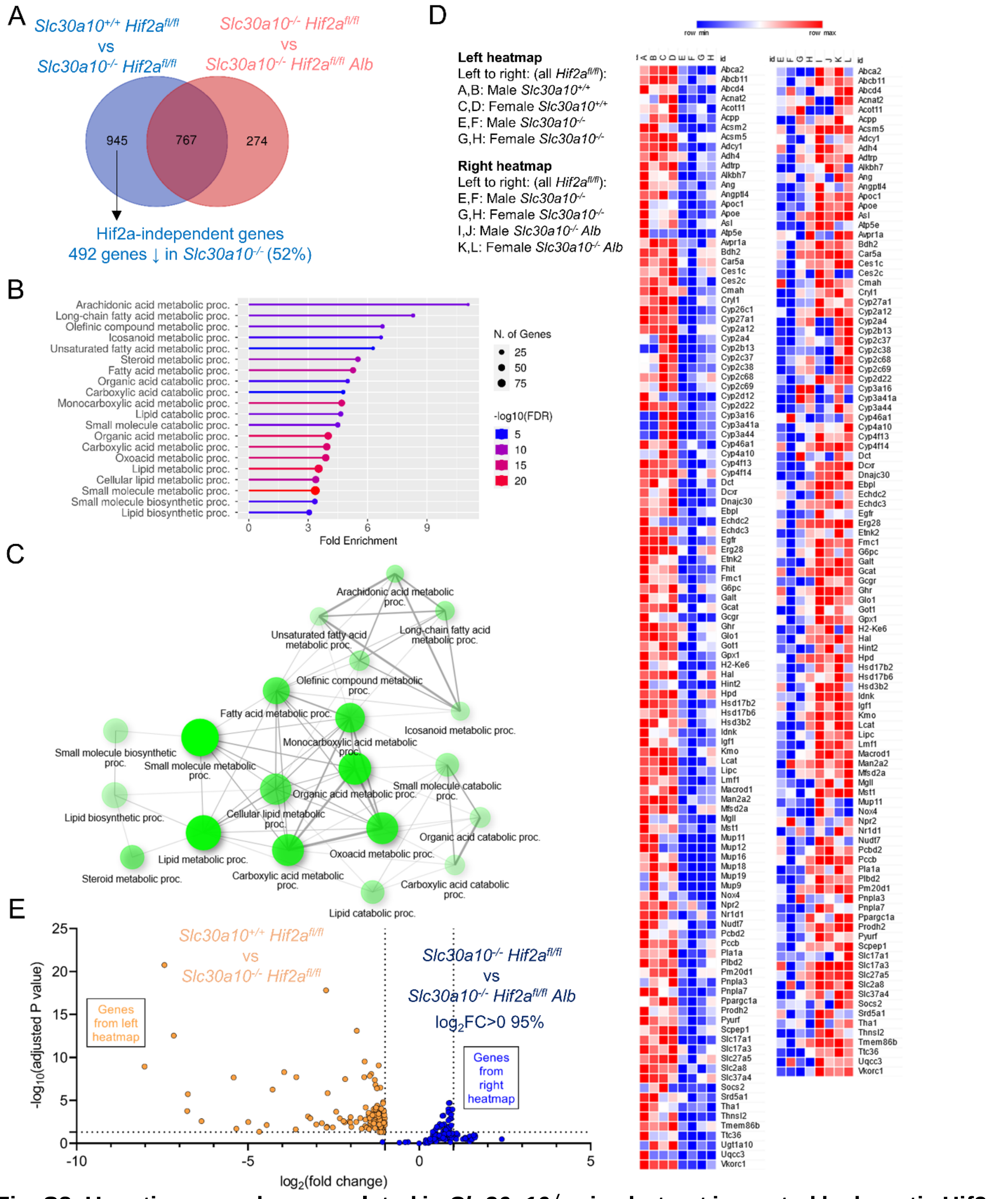


**Fig. S8: Hepatic genes downregulated in *Slc30a10*$^{-/-}$ mice but not impacted by hepatic Hif2a deficiency align largely with metabolic processes.** (**A**) Venn diagram partially reproduced from Fig. 8G. (**B, C**) Gene enrichment analysis. (**D**) Heat map of differentially expressed genes aligned with pathways encircled by dashed line in (**C**). (**E**) Volcano plot of genes from left heatmap from (**D**) plotted as orange circles and from right heatmap from (**D**) as blue circles, indicating that 95% genes not significantly impacted by hepatic Hif2a deficiency are upregulated, i.e. $\log_2$(fold change)>0.

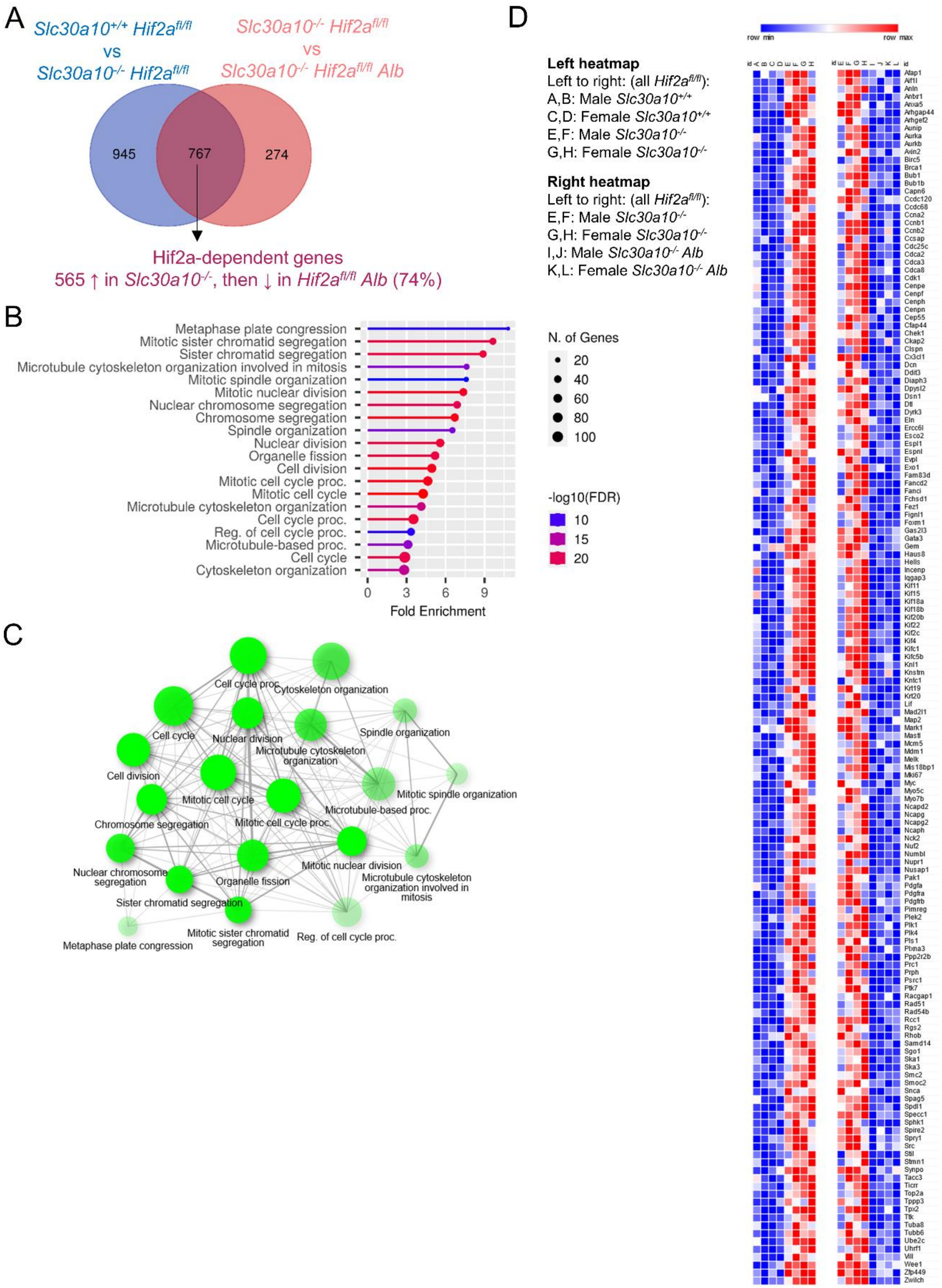


**Fig. S9: Hepatic genes upregulated in *Slc30a10*$^{-/-}$ mice then downregulated with hepatic Hif2a deficiency align with cell cycle.** (**A**) Venn diagram partially reproduced from Fig. 8G. (**B, C**) Gene enrichment analysis. (**D**) Heat map of differentially expressed genes aligned with pathway shown in (**C**).

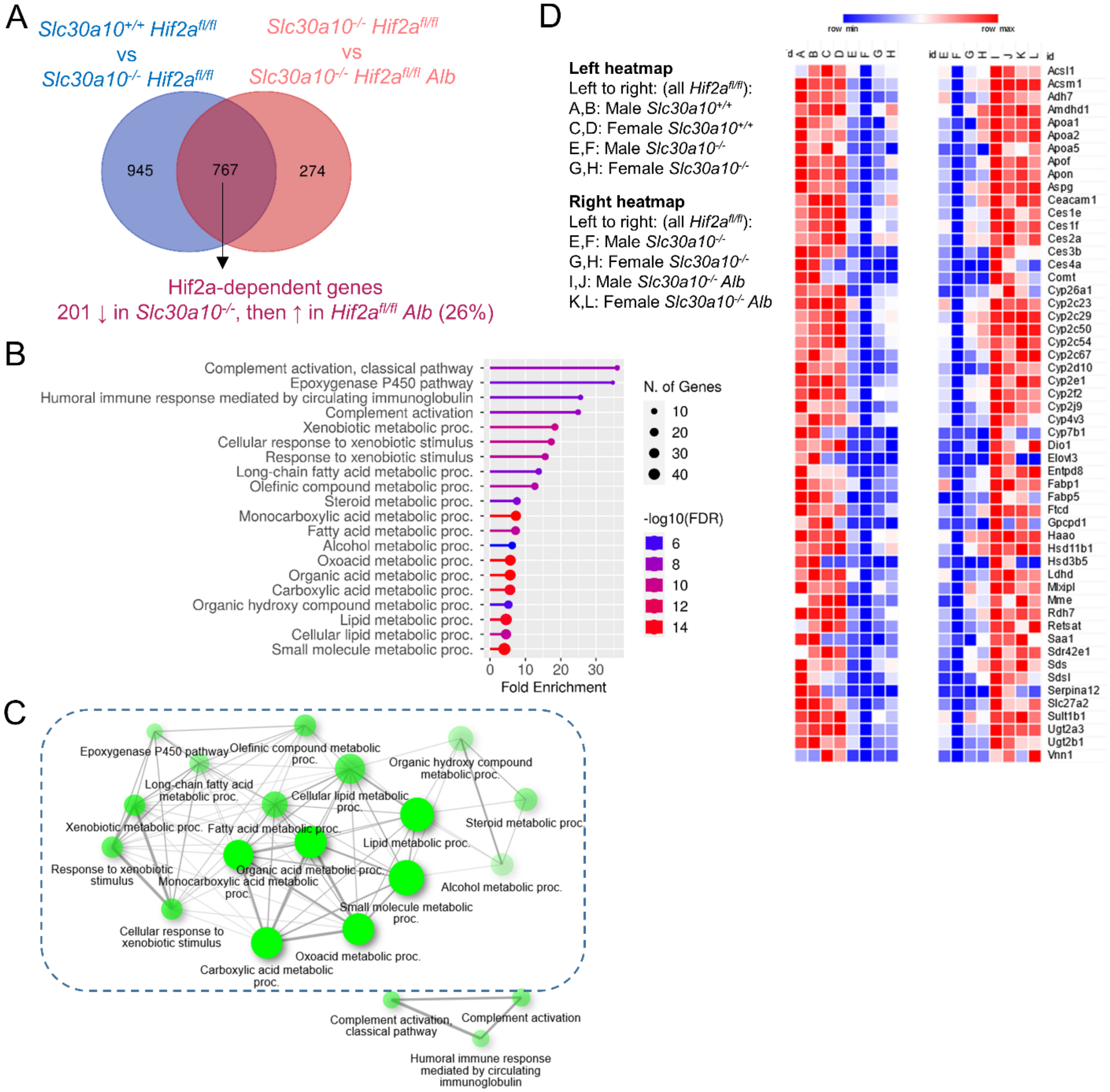


**Fig. S10: Hepatic genes downregulated in *Slc30a10*$^{-/-}$ mice then upregulated with hepatic Hif2a deficiency align largely with metabolic processes.** (**A**) Venn diagram partially reproduced from Fig. 8G. (**B, C**) Gene enrichment analysis. (**D**) Heat map of differentially expressed genes aligned with pathway encircled by dashed line in (**C**).

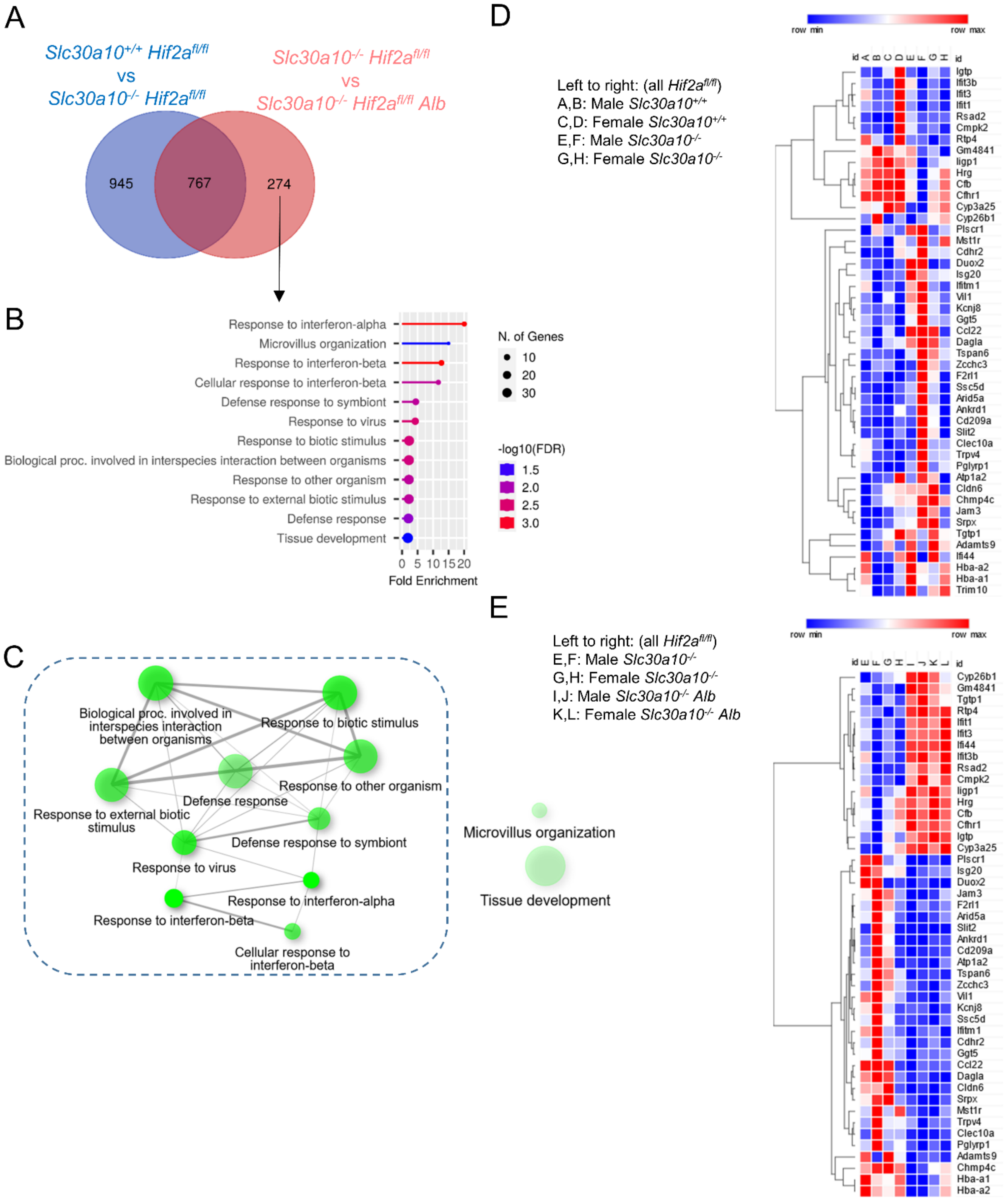


**Fig. S11: Hepatic genes not downregulated in *Slc30a10*$^{-/-}$ mice but differentially expressed with hepatic Hif2a deficiency align largely with stimuli response pathways.** (**A**) Venn diagram partially reproduced from Fig. 8G. (**B, C**) Gene enrichment analysis. (**D, E**) Heat map of gene expression in *Slc30a10*$^{-/-}$ *Hif2a*$^{fl/fl}$ vs. *Slc30a10*$^{+/+}$ *Hif2a*$^{fl/fl}$ mice (**D**) and in *Slc30a10*$^{-/-}$ *Hif2a*$^{fl/fl}$ *Alb* vs. *Slc30a10*$^{-/-}$ *Hif2a*$^{fl/fl}$ mice (**E**) for genes aligned with pathways encircled by dashed shape in (**C**).

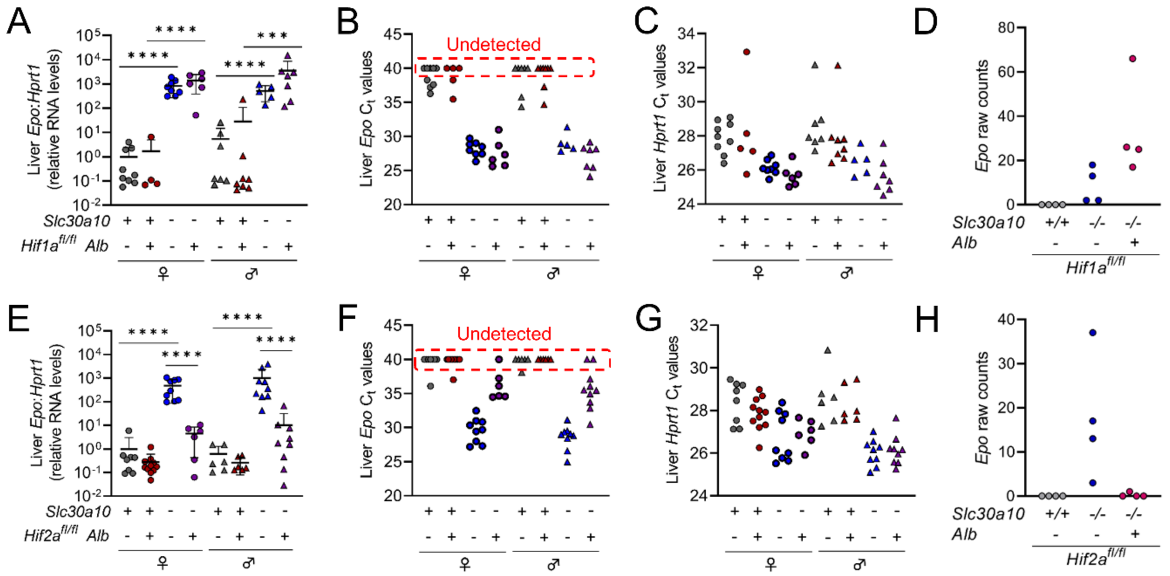


**Fig. S12: qPCR $C_t$ values and RNA-seq raw counts for *Epo*.** (**A-D**) Reproduction of liver *Epo* RNA levels from Fig. 4F (**A**), along with *Epo* (**B**) and *Hprt1* (**C**) Ct values from qPCR and *Epo* raw counts from RNA-seq (**D**). (**E-H**) Reproduction of liver *Epo* RNA levels from Fig. 5F (**E**), along with *Epo* (**F**) and *Hprt1* (**G**) Ct values from qPCR and *Epo* raw counts from RNA-seq (**H**).
